# Inhibition of SF3B1 affects recruitment of P-TEFb to chromatin through multiple mechanisms

**DOI:** 10.1101/2024.06.26.600844

**Authors:** Gilbert Ansa, Shona Murphy, Michael Tellier

## Abstract

Processing of nascent pre-mRNAs is tightly coupled to transcription by RNA polymerase II (RNAPII) through reversible phosphorylation of the polymerase and associated factors by transcriptional kinases. P-TEFb, comprising cyclin-dependent kinase (CDK)9 and cyclin T1, is a key transcription elongation kinase, which also regulates co-transcriptional splicing and mRNA cleavage and polyadenylation. Chemical inhibition of SF3B1, a component of the splicing factor U2 snRNP, decreases P-TEFb recruitment to chromatin and mirrors the effect of P-TEFb inhibition on transcription. However, the mechanism of this effect of SF3B1 inhibitors was unclear. Here we show that SF3B1 inhibition causes rapid nuclear export of P-TEFb and loss of SF3B1 phosphorylation. SF3B1 is in complex with P-TEFb on chromatin with the elongation/splicing factor HTATSF1 and the splicing factor SNW1. SF3B1 inhibition causes the nuclear export of SNW1, but not of HTATSF1. The chromatin association of AFF4, an interaction partner of P-TEFb, is also affected by SF3B1 inhibition. Surprisingly, SF3B1 inhibition promotes degradation of SRSF2, a splicing factor known to help recruit P-TEFb to chromatin. Our results indicate that SF3B1 inhibition affects P-TEFb recruitment to genes via multiple pathways. Together, these interactions ensure efficient coupling of transcription and splicing.

## INTRODUCTION

The expression of a mature mRNA from an intron-containing protein-coding gene requires transcription by RNA polymerase II (RNAPII) and co-transcriptional RNA processing, including mRNA capping, pre-mRNA splicing, and mRNA cleavage and polyadenylation. The C-terminal domain (CTD) of RNAPII functions as a recruiting platform for RNA processing factors (1,2) to physically and functionally link transcription and RNA processing (3,4). Post-translational modifications of the RNAPII CTD heptapeptide (Tyr1Ser2Pro3Thr4Ser5Pro6Ser7) by multiple cyclin-dependent kinases (CDKs) during transcription, especially phosphorylation of Ser2 and Ser5 residues (5), helps to recruit capping, splicing, and cleavage and polyadenylation factors (3,4).

Following transcription initiation, binding of negative elongation factor (NELF) and of 5,6-dichloro-1-b-D-ribofuranosylbenzimidazole (DRB)-sensitivity inducing factor (DSIF) complexes to RNAPII promotes its pausing 20-40 nucleotides downstream of the transcription start site (TSS) (6,7). Positive transcription elongation factor b (P-TEFb) is a complex of CDK9 and Cyclin T1 that regulates RNAPII pause release via phosphorylation of the RNAPII CTD and of two negative elongation complexes, NELF and DSIF, resulting in the release of NELF from chromatin and the transformation of DSIF into a positive elongation factor (8). Recruitment of P-TEFb to RNAPII can occur via multiple pathways (9), including factors involved in transcription such as BRD4 (10,11), MYC (12,13), and AFF4 as part of the super elongation complex (SEC) (14,15), but also RNA-binding and splicing factors including DDX21 (16), HTATSF1 (17,18), RBM7 (19), RBM22 (20), SNW1 (21,22), SFPQ (23), SRSF2 (24,25), and WDR43 (26).

Chemical inhibition of SF3B1, a key member of the U2 snRNP complex which recognises the branch point during splicing (27), by the small molecules Spliceostatin A (SSA) (28) and Pladienolide B (PlaB) (29) reduces the recruitment of Cyclin T1 to chromatin (30) and promotes increased RNAPII pausing, decreased transcription elongation, and causes premature termination of transcription via increased intronic poly(A) site usage (30-35), presumably through loss of P-TEFb kinase activity.

SF3B1 is a protein that contains more than 50 phosphorylation sites in its N-terminus, upstream of its HEAT domain where cancer-associated mutations affecting 3’ splice site recognition are found (36). SF3B1 is phosphorylated by multiple kinases, including CDK9 (37,38), but also CDK1 (39), CDK2 (39,40), CDK7 (41), CDK11 (42), CDK12 (43), and DYRK1A (44,45), whereas protein phosphatase (PP)1 and PP2A have been found to dephosphorylate SF3B1 (38,46-48). SF3B1 phosphorylation is dynamically regulated during pre-mRNA splicing, with phosphorylation occurring during spliceosome activation (49-52) followed by its dephosphorylation after the second catalytic step of splicing (46,49,52). SF3B1 phosphorylation also regulates its interaction with NIPP1 (53), a protein recruiting PP1 to dephosphorylate SF3B1 (48). SF3B1 phosphorylation is also dynamic during the cell cycle (39), with a peak in phosphorylation during the G1/S phase, which correlates with increased association of SF3B1 with chromatin (39,52,54).

The mechanism linking SF3B1 inhibition to the loss of P-TEFb recruitment to chromatin was unclear. In this study, we have investigated what happens to P-TEFb and SF3B1 phosphorylation following SF3B1 inhibition with PlaB (29) and the splicing inhibitor Herboxidiene (HB) (55). Mechanistically, SF3B1 is in complex on chromatin with HTATSF1, SNW1, and P-TEFb. PlaB treatment causes export of P-TEFb and SNW1 to the cytoplasm. A complex of SF3B1, HTATSF1, and P-TEFb remains on chromatin after SF3B1 inhibition but is not sufficient to overcome the loss of other interactions in promoting RNAPII pause release and SF3B1 phosphorylation. Two other factors known to recruit P-TEFb to chromatin for RNAPII pause release, AFF4 and SRSF2, are also affected by SF3B1 inhibition; AFF4 is lost from chromatin and SRSF2 is rapidly degraded by the nuclear proteasome, which would further contribute to the loss of P-TEFb from chromatin. Additionally, we show that P-TEFb and CDK12 phosphorylate several residues of SF3B1.

Overall, our data shows that SF3B1 inhibition affects P-TEFb recruitment via multiple pathways and confirms that SF3B1 is a target of both CDK9 and CDK12. Interaction between SF3B1 and P-TEFb and phosphorylation of SF3B1 by CDK9 and CDK12 therefore ensure that productive elongation is efficiently coupled to productive splicing of pre-mRNA.

## MATERIAL AND METHODS

### Cell culture

HeLa and HEK293 wild-type were obtained from ATCC (ATCC® CCL-2™ and ATCC® CRL-1573™, respectively) while HEK293 CDK9^as^ cells are from (38). The three cell lines were grown in DMEM medium supplemented with 10% foetal bovine serum, 100 U/ml penicillin, 100 µg/ml streptomycin, and 2 mM L-glutamine at 37°C and 5% CO_2_. HeLa or HEK293 wild-type cells were treated with 100 nM or 1 µM of Pladienolide B (PlaB, Cayman Chemical Company), 1 µM of Herboxidiene (HB, Tebubio), 100 µM of 5,6-dichlorobenzimidazone-1-β-D-ribofuranoside (DRB, Sigma), 0.02 µM of Leptomycin B (LMB, Cell Signaling Technology), 1 µM of Bortezomib (Bort, Cayman Chemical), or 10 ng/ml of TNFα (Peprotech) for the time indicated in the figures. HEK293 CDK9^as^ cells were treated with 15 µM 1-NA-PP1 (Cayman Chemical Company) for 30 min. As a negative control, HeLa, HEK293 wild-type, or HEK293 CDK9^as^ cells were treated with DMSO, the resuspension vehicle of PlaB, HB, DRB, Bortezomib, and 1-NA-PP1, with ethanol, the resuspension vehicle of LMB, or with water, the resuspension vehicle of TNFα. Cells were routinely checked for mycoplasma contamination using Plasmo Test Mycoplasma Detection Kit (InvivoGen, rep-pt1).

### RNA purification, RT-PCR, and qRT-PCR

RNA was extracted from ∼70% confluent HeLa cells grown in 6-wells plate using the RNeasy Micro Kit (Qiagen) with a DNase step according to the manufacturer’s instruction. Reverse transcription was performed with 500 ng of RNA using random hexamers or oligo(dT) 12-18 primer with the SuperScript III kit (Invitrogen) according to the manufacturer’s instruction. PCR reactions were performed with the Phusion High Fidelity DNA polymerase (NEB) according to the manufacturer’s instruction. Quantifications were performed with Image Studio Lite version 5.2.5 and shown as % of unspliced RNA ((unspliced RNA signal / (spliced RNA signal + unspliced RNA signal) * 100). For qRT-PCR, cDNA was amplified with a QuantiTect SYBR Green PCR kit (QIAGEN) and a Rotor-Gene RG-3000 (Corbett Research). Values are normalised to the 7SK non-coding RNA (random hexamers) or the GAPDH mRNA (oligo(dT) primer) that are used as internal controls. Experiments were performed in technical triplicates and to ensure reproducibility, as biological duplicates or triplicates as indicated on the Figures. Primers used for RT-PCR and qRT-PCR are shown in Table 1.

**Table 1.**
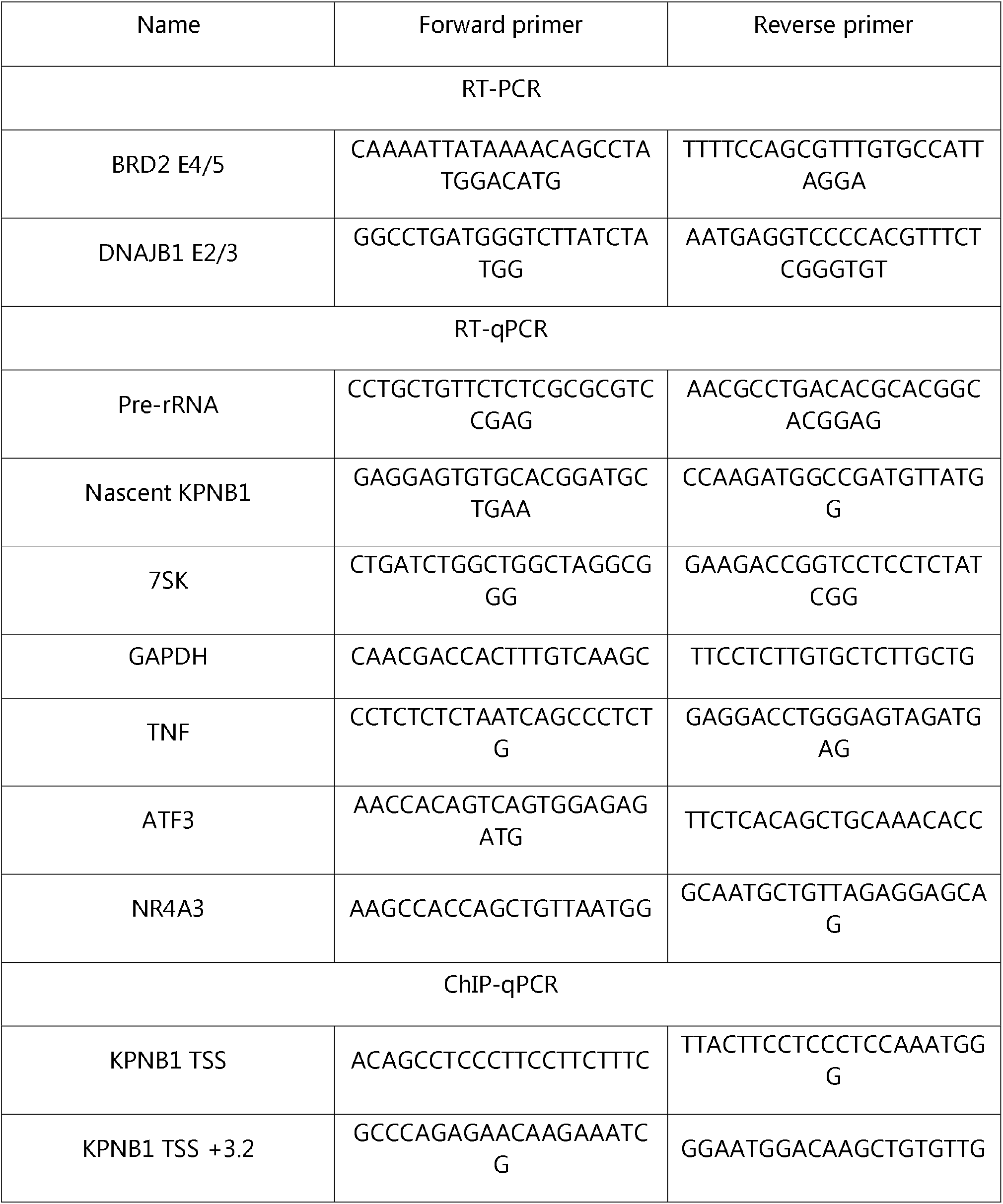
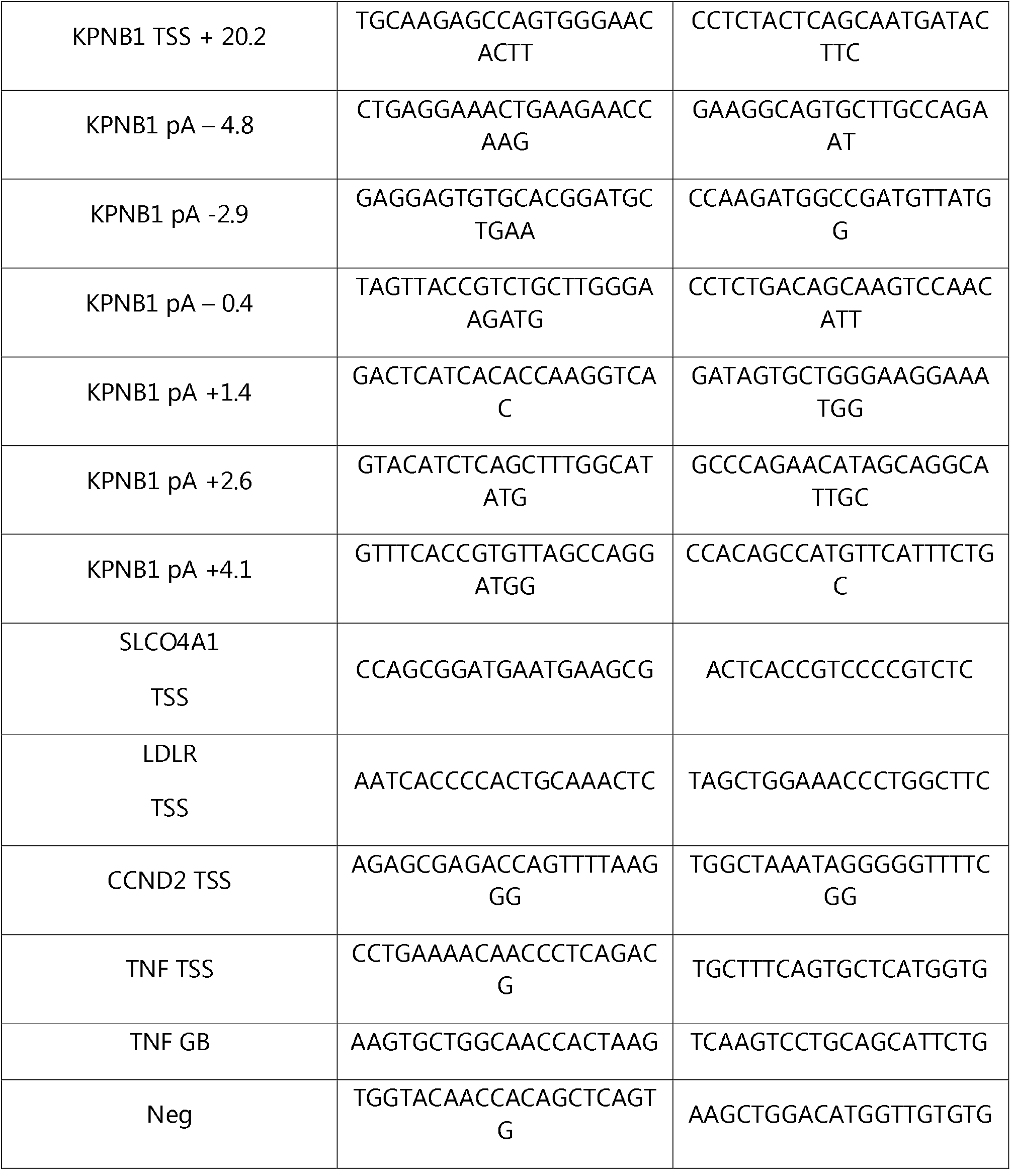
Primers used in this study.

### ChIP-qPCR

ChIP were performed as previously described (56). HeLa cells were grown in 150 mm dishes until ∼80% confluency. The cells were fixed with 1% formaldehyde for 10 min at room temperature on a shaker before being quenched with 125 mM glycine for 5 min at room temperature with shaking. The cells were washed twice with ice-cold PBS, scraped with ice-cold PBS, and pelleted at 1,500 g for 5 min at 4°C. Cell pellets were resuspended in ChIP lysis buffer (10□mM Tris-HCl pH 8.0, 0.25% Triton X-100, 10□mM EDTA, protease inhibitor cocktail, and phosphatase inhibitor), incubated on ice for 10 min, and centrifuged at 1,500 g for 5 min at 4°C. Pellets were resuspended in ChIP wash buffer (10□mM Tris–HCl pH 8.0, 200□mM NaCl, 1□mM EDTA, protease inhibitor cocktail, and phosphatase inhibitor) and centrifuged at 1,500 g for 5 min at 4°C. Pellets were then resuspended in ChIP sonication buffer (10□mM Tris–HCl pH 8.0, 100□mM NaCl, 1□mM EDTA, protease inhibitor cocktail, and phosphatase inhibitor), incubated on ice for 10 min, and sonicated for 30□cycles, 30 s on/30 s off using a Bioruptor Pico (Diagenode). Chromatin was pelleted at 15,800 g for 15 min at 4°C. The supernatant was precleared for on a rotating wheel at 16 rpm 30 min at 4°C with 10 µl of Protein G dynabeads (Thermo Fisher) that were previously washed with 100 µl of RIPA buffer (10□mM Tris–HCl pH 8.0, 150□mM NaCl, 1□mM EDTA, 0.1% SDS, 1% Triton X-100, 0.1% sodium deoxycholate). Following chromatin quantification with a Nanodrop One, 60 – 100 µg of chromatin was incubated overnight on a rotating wheel at 16 rpm at 4°C with the antibody of interest (Table 2). For each IP, 15 µl of Protein G dynabeads were washed with 150 µl of RIPA buffer and incubated overnight on a rotating wheel at 16 rpm at 4°C with 15 µl of RIPA containing 4 mg/ml of bovine serum albumin (BSA). Pre-blocked protein G dynabeads and chromatin were mixed for 1 h on a rotating wheel at 16 rpm at 4°C before being washed three times with 300 µl ice-cold RIPA, three times with 300 µl ice-cold High Salt Wash Buffer (10□mM Tris–HCl pH 8.0, 500□mM NaCl, 1□mM EDTA, 0.1% SDS, 1% Triton X-100, 0.1% sodium deoxycholate), twice with 300 µl ice-cold LiCl Wash Buffer (10□mM Tris-HCl pH 8.0, 250□mM LiCl, 1□mM EDTA, 1% NP-40, 1% sodium deoxycholate), and twice with 300 µl ice-cold TE buffer (10□mM Tris-HCl pH 7.5, 1□mM EDTA). Each sample was eluted twice from the protein G dynabeads with 50□μl of Elution buffer (100□mM NaHCO3, 1% SDS, 10□mM DTT) on a Thermomixer for 15□min at 25°C at 1,400□rpm. For each input sample, which corresponds to the supernatant of the IgG condition after mixing chromatin and protein G dynabeads), 90□μl of Elution buffer was added to 10□μl total input. Each sample was treated with RNase A (0.6□μl of 10□mg/ml) for 30□min at 37°C followed by the addition of 200□mM NaCl and incubation at 65°C for 5 h to reverse the crosslinks. Precipitation was performed overnight at ‐20°C following the addition of 2.5x volume of 100% ethanol. Ethanol was removed after a 20 min centrifugation at 15,800□g at 4°C and pellets resuspended in 100□μl TE, 25□μl 5x Proteinase K buffer (50□mM Tris–HCl pH 7.5, 25□mM EDTA, 1.25% SDS) and 1.5□μl Proteinase K (20□mg/ml). The samples were incubated 2□h at 45°C and DNA purified with a Qiagen PCR Purification Kit before being kept at ‐20°C. qPCRs were performed with a QuantiTect SYBR Green PCR kit (QIAGEN) and a Rotor-Gene RG-3000 (Corbett Research). Signals are presented as (IP – IgG) / Input. Experiments were performed in technical triplicates and to ensure reproducibility, as biological replicates as indicated on the respective Figures. Primers used for ChIP-qPCR are shown in Table 1.

### 5’EU

HeLa cells were seeded at 40-50% confluency on coverslips in a 6-wells plate and treated the following day with DRB or PlaB for the time indicated on the figure before being incubated with 1 mM 5-ethylnyl uridine (5-EU) for 1 h to label newly synthesized RNA. Cells were then fixed by transferring coverslips to 3.7 % formaldehyde diluted in PBS for 15 min, washed with PBS, and then permeabilised in 0.5 % Triton X-100 diluted in PBS for 15 min at room temperature. Global nascent RNA transcription was detected using Click-iT® RNA Alexa Fluor 488 Imaging kit (ThermoFisher, C10329) according to the manufacturer’s instructions. Following Click-iT reaction, cellular DNA was immunostained with Hoechst 33342 diluted 1:1000 in PBS for 15 min at room temperature, protected from light. Coverslips were then washed twice with PBS, then mounted onto slides using Fluoromount-G (SouthernBiotech, 0100-01) containing DAPI. Slides were visualised with an inverted multi-channel fluorescence microscope (Evos M7000; ThermoFisher). Fluorescent intensity was analysed and quantified with ImageJ (version 1.53q) by first using the DAPI channel (λ_Ex_/λ_Em_ (with DNA) = 350/461 nm) image to define the nucleus, then GFP channel (λ_Ex_/λ_Em_ = 495/519 nm) to measure the fluorescent intensity within the nucleus. Data were plotted by box-and-whisker plot with GraphPad Prism 9.5 software with the following settings: boxes: 25-75 percentile range; whiskers: min-max values; horizontal bars: median.

### Immunofluorescence

HeLa cells were seeded on 13 mm sterile round coverslips overnight at 37°C to get 60-70% confluent cells. Following respective drug treatments, cells on coverslips were washed twice with PBS then fixed using 4% paraformaldehyde (Sigma Aldrich) for 15 min at room temperature. Coverslips were washed twice with PBS then permeabilised and treated with blocking solution (0.1% Triton X-100, 10% FBS, and PBS) for 30 min at room temperature. Blocking solution was removed and coverslips were incubated with primary antibodies (Rabbit anti-G3BP1 (17798, Cell Signaling), 1:500; Mouse anti-CDK9 (D-7) (sc-13130, Santa Cruz Biotechnology), 1:500; Rabbit anti-SC35 (SRSF2) (ab204916, Abcam), 1:100) diluted in blocking solution overnight at 4°C. Following incubation, primary antibody solution was removed, and coverslips washed three times with PBS for 5 min each. Coverslips were then incubated with secondary antibody (Goat anti-Mouse IgG (H+L) Secondary Antibody, DyLight™ 488 (35502, Invitrogen), 1:500; Goat anti-Rabbit IgG (H+L) Highly Cross-Abssorbed Secondary Antibody, Alexa Fluor™ Plus 488 (A-11034, Invitrogen), 1:500) diluted in solution (1% FBS and PBS) for 1 h at room temperature. Following incubation, secondary antibody solution was removed, and coverslips washed three times with PBS for 5 min each. Coverslips were incubated in Hoechst 33342 solution (62249, Thermo Fisher) (diluted 1:1,000 in PBS) protected from light for 10 min at room temperature. Hoechst 33342 solution was removed then coverslips were washed twice with PBS and mounted upside down on a drop of FluorSave Reagent (345789, Millipore) on a clean glass slide. Coverslips were sealed to glass slides using nail polish then left overnight, protected from light at room temperature to dry. Once dry, glass slides were stored at 4°C, protected from light until imaging. Cells were detected and imaged by a Zeiss 880 Airyscan microscope. Antibody validation is provided on the manufacturer’s website.

### Cell fractionation

For cytoplasmic fractions, HeLa cells were washed twice with cold PBS then scraped into 15 ml tubes and centrifuged at 1,000 rpm for 5 min at 4°C. The pellet was resuspended thoroughly in 1 ml of Lysis buffer B (10 mM Tris-HCl (pH 8-8.4), 140 mM NaCl 1.5 mM MgCl2, and 0.5% NP40) supplemented with protease inhibitor cocktail (Roche) and PhosSTOP (Roche) by slow pipetting and then transferred to a 1.7 ml tube. The lysate was centrifuged at 1,000 g for 3 min at 4°C then 500 μl of the supernatant of the unpurified cytoplasmic fraction was transferred to a new tube. Purified cytoplasmic fraction was achieved by centrifugation at 10,000 g for 1 min at 4°C.

For nucleoplasm and chromatin fractions, the rest of the unpurified cytoplasmic fraction was discarded and the nuclear pellet resuspended in 1 ml of Lysis buffer B. Following the transfer to a 14 ml round bottom Falcon tube, 100 µl of Detergent stock (3.3% (w/v) Sodium Deoxycholate, 6.6% (v/v) Tween 40) was added drop-by-drop under slow vortexing. After a 5 min incubation on ice, the suspension was transferred to a new ice-cold 1.7 ml tube and centrifuged a 1,000 g for 3 min at 4°C. The nuclear pellet was resuspended in 1 ml of Lysis buffer B. Following centrifugation at 1,000 g for 3 min at 4°C, the nuclear pellets were resuspended by pipetting up and down in 125 μl of NUN1 buffer (20 mM Tris–HCl pH 7.9, 75 mM NaCl, 0.5 mM EDTA and 50% (vol/vol) glycerol) and moved to a new 1.5 ml ice-cold microcentrifuge tube. Following the addition of 1.2 ml of ice-cold NUN2 buffer (20 mM HEPES-KOH pH 7.6, 300 mM NaCl, 0.2 mM EDTA, 7.5 mM MgCl_2_, 1% (vol/vol) NP-40 and 1 M urea), the tubes were vortexed at maximum speed for 10 s and incubated on ice for 15 min with a vortexing step of 10 s every 3 min. The samples were centrifuged at 16,000 g for 10 min at 4°C and the supernatant kept at the nucleoplasm fraction while the chromatin pellets were washed with 500 μl of ice-cold PBS and then with 100 μl of ice-cold water. The chromatin pellet was then digested in 100 μl of water supplemented with 1 μl of Benzonase (25–29 units, Merck Millipore) for 15 min at 37°C in a thermomixer at 1,400 rpm.

For direct purification of the nucleoplasm and chromatin fraction, a ∼ 80% confluent 15 cm dish was washed twice with ice-cold PBS and scrapped in 5 ml of ice-cold PBS. The cells were pelleted at 420 g for 5 min at 4°C. After discarding the supernatant, the cells were resuspended in 4 ml of ice-cold HLB+N buffer (10 mM Tris–HCl pH 7.5, 10 mM NaCl, 2.5 mM MgCl_2_ and 0.5% (vol/vol) NP-40) and incubated on ice for 5 min. The cell pellets were then underlayed with 1 ml of ice-cold HLB+NS buffer (10 mM Tris–HCl pH 7.5, 10 mM NaCl, 2.5 mM MgCl_2_, 0.5% (vol/vol) NP-40 and 10% (wt/vol) sucrose). Following centrifugation at 420 g for 5 min at 4°C, the nuclear pellets were resuspended by pipetting up and down in 125 μl of NUN1 buffer and processed as described above

### Co-immunoprecipitation

For whole-cell co-immunoprecipitation, we prepared for each sample and IgG control: 120□μl of Dynabeads M-280 Sheep anti-mouse IgG (Thermo Fisher) or 40□μl of Dynabeads protein G (Thermo Fisher) were pre-blocked overnight at 4°C on a wheel at 16□rpm in 1□ml of PBS supplemented with 0.5% BSA. The next day, the beads were washed three times in IP buffer (25□mM Tris–HCl pH 8.0, 150□mM NaCl, 0.5% NP-40, 10% Glycerol, 2.5□mM MgCl_2_), before being incubated for 2□h at 4°C on a wheel at 16□rpm in 600□μl of IP buffer supplemented with 5□μg of SF3B1 antibody (D221-3, MBL International), 5□μg of Cyclin T1 antibody (81464, Cell Signaling), 5□μg of CDK9 antibody (sc-13130, Santa Cruz Biotechnology), 5□μg of HTATSF1 antibody (A302-023A, Bethyl Laboratories), or Normal Rabbit IgG (2729S, Cell Signaling Technology) and protease inhibitor cocktail (cOmplete™ EDTA-free Protease Inhibitor Cocktail, Sigma-Aldrich). In the meantime, a 70–80% confluent 15□cm dish of HeLa cells was washed twice with ice-cold PBS and scrapped with ice-cold PBS supplemented with protease inhibitor cocktail. The cells were pelleted at 500□g for 5□min at 4°C. The pellets were resuspended in 800□μl of Lysis buffer (50□mM Tris-HCl pH 8.0, 150□mM NaCl, 1% NP-40, 10% glycerol, 2.5□mM MgCl_2_, protease inhibitor cocktail, PhosSTOP (Sigma-Aldrich), 1× PMSF (Sigma-Aldrich), and 25–29□units of Benzonase (Merck Millipore)) and incubated at 4°C on a wheel at 16□rpm for 30□min. After centrifuging for 15□min at 13,000□g at 4°C, 800□μl of Dilution buffer (150□mM NaCl, 10% glycerol, 2.5□mM MgCl_2_, protease inhibitor cocktail, PhosSTOP, and 1× PMSF) was added to each supernatant.

The beads conjugated with antibodies were washed three times with IP buffer supplemented with protease inhibitor cocktail before being incubated with 1□mg of proteins at 4°C on a wheel at 16□rpm for 2□h. The beads were washed three times with IP buffer supplemented with protease inhibitor cocktail and three times with IP buffer without NP-40 supplemented with protease inhibitor cocktail. Proteins were eluted in 40□μl of 1× LDS plus 100□mM DTT for 10□min at 70°C.

For co-immunoprecipitation experiments performed on cellular fractionation, rather than using 1 mg of whole cell proteins, 1 mg of proteins purified as described above were used for each fraction.

### Western blotting

For whole-cell extract, cells were washed twice in ice-cold PBS and collected in ice-cold PBS by centrifugation at 800 g for 5 min at 4°C. The pellets were resuspended in RIPA buffer supplemented with protease inhibitor cocktail and PhosSTOP, kept on ice for 30 min with a vortexing step every 10 min. After centrifugation at 14,000 g for 15 min at 4°C, the supernatants were kept and quantified with the Bradford method. For loading, 20 μg of proteins were boiled in 1× LDS plus 100 mM DTT.

Western blots were performed with NuPAGE Novex 4–12% Bis–Tris Protein Gels (Life Technologies) with the following primary antibodies: Rpb1 NTD (D8L4Y) Rabbit mAb (14958S, Cell Signaling Technology), Phospho-Rpb1 CTD (Ser2) (E1Z3G) Rabbit mAb (13499S, Cell Signaling Technology), Rabbit anti-SF3b155/SAP155 (A300-996A, Bethyl Laboratories), Rabbit anti-Phospho-SF3B1 Ser129 (PD043, MBL International), Rabbit anti-Phospho-SF3B1 Thr142 ((38)), Rabbit anti-Phospho-SF3B1 Thr211 (PA5-105427, Invitrogen), Rabbit anti-Phospho-SF3B1 Thr313 (D8D8V, Cell Signaling), Rabbit anti-Tat-SF1 (A302-023A, Bethyl Laboratories), Rabbit anti-CDK9 (ab76320, Abcam), Rabbit anti-Phospho-CDK9 (pThr186) (2549, Cell Signaling), Rabbit anti-Cyclin T1 (ab184703, Abcam), Rabbit anti-SNW1 (A300-784A, Bethyl Laboratories), Rabbit anti-c-MYC (sc-764, Santa Cruz Biotechnology), Rabbit anti-SC35 (SRSF2) (ab204916, Abcam), Rabbit anti-BRD4 (A301-985A100, Bethyl Laboratories), Rabbit anti-MCEF (AFF4) (A302-539A, Bethyl Laboratories), Mouse anti-U1A (3F9-1F7, Novus Biologicals), Rabbit anti-Actin (C-11) (sc-1615, Santa Cruz Biotechnology), Rabbit anti-α-Tubulin (2144S, Cell Signaling), Rabbit anti-Nucleolin (ab305947), Abcam), and Histone H3 (ab1791, Abcam). Secondary antibodies were purchased from Merck Millipore (Goat Anti-Rabbit IgG Antibody, HRP-conjugate, 12-348, and Goat Anti-Mouse IgG Antibody, HRP conjugate, 12-349), the chemiluminescent substrate (SuperSignal West Pico PLUS) from Thermo Fisher, and the membranes visualized on an iBright FL1000 Imaging System (Thermo Fisher).

### mNET-seq

mNET-seq was carried out as previously described in (56). In brief, the chromatin fraction was isolated from 4 × 10^7^ HeLa cells treated with DMSO or PlaB for 30 min. Chromatin was digested in 100 μl of MNase (40 units/μl) reaction buffer for 2 min at 37°C in a thermomixer at 1,400 rpm. MNase was inactivated with the addition of 10 μl EGTA (25mM) and soluble digested chromatin was collected after centrifugation at 13 000 rpm for 5 min at 4°C. The supernatant was diluted with 400 μl of NET-2 buffer and antibody-conjugated beads were added. Antibodies used: Pol II (MABI0601, MBL International), Ser2P (MABI0602, MBL International) and Ser5P (MABI0603, MBL International). Immunoprecipitation was performed at 4°C on a rotating wheel at 16 rpm for 1 h. The beads were washed six times with 1 ml of ice-cold NET-2 buffer and once with 100 μl of 1x PNKT (1x PNK buffer and 0.05% Triton X-100) buffer. Washed beads were incubated in 200 μl PNK reaction mix for 6 min in a Thermomixer at 1,400 rpm at 37°C. The beads were then washed once with 1 ml of NET-2 buffer and RNA was extracted with Trizol reagent. RNA was suspended in urea Dye and resolved on a 6% TBU gel (Invitrogen) at 200 V for 5 min. In order to size select 35–100 nt RNAs, a gel fragment was cut between BPB and XC dye markers. For each sample, several small holes were made with a 25G needle at the bottom of a 0.5 ml tube, which is then placed in a 1.5 ml tube. Gel fragments were placed in the layered tube and broken down by centrifugation at 12,000 rpm for 1 min. The small RNAs were eluted from the gel using RNA elution buffer (1 M NaOAc and 1 mM EDTA) for 1 h on a rotating wheel at 16 rpm at room temperature. Eluted RNAs were purified with SpinX column (Coster) and two glass filters (Millipore) and the resulting flow-through RNA was precipitated overnight with ethanol. RNA libraries were prepared according to manual of Truseq small RNA library prep kit (Illumina). Deep sequencing (Hiseq4000, Illumina) was conducted by the high throughput genomics team of the Wellcome Trust Centre for Human Genetics (WTCHG), Oxford.

### Gene annotation

The list of protein-coding genes, intronless genes, and histone genes was extracted from the Gencode V35 annotation as in (57). Briefly, a list of 9,883 expressed protein-coding genes was obtained by applying a 2□kb size cut-off. The TSS and poly(A) site of each gene was defined by the highest transcript isoform expressed in two nuclear poly(A)+ RNA-seq in HeLa cells (58) at more than 0.1 transcript per million (TPM), following quantification of transcript expression with Salmon version 1.10.2 (59). The list of used exons was obtained by extracting the location of exons from each most expressed transcript of the 9,883 protein-coding genes from Gencode V35. The list of expressed intronless and histone genes was obtained by extracting expressed protein-coding genes with only one exon and then separating them based on histone and non-histone genes. For K562 gene expression level, biological replicates of RNA-seq treated with DMSO were obtained from GSE148433 (30) and quantified with Salmon, keeping the highest protein-coding gene transcript expressed per gene.

### mNET-seq analysis

Total RNAPII mNET-seq in Raji CDK9as cells treated with DMSO or 1-NA-PP1 were obtained from GSE96056 (60). Adapters were trimmed with Cutadapt version 1.18 (61) in paired-end mode with the options: --minimum-length 10 -q 15,10 -j 16 -A GATCGTCGGACTGTAGAACTCTGAAC -a AGATCGGAAGAGCACACGTCTGAACTCCAGTCAC. Trimmed reads were mapped to the human GRCh38.p13 reference sequence with□STAR version 2.7.3a (62) and the parameters: --runThreadN 16 --readFilesCommand gunzip -c -k -- limitBAMsortRAM 20,000,000,000 --outSAMtype BAM SortedByCoordinate. SAMtools version 1.9 (63) was used to keep the properly paired and mapped reads (-f 3). A custom python script (32) as used to obtain the 3’ nucleotide of the second read and the strandedness of the first read. Strand-specific bam files were generated with SAMtools. FPKM-normalised bigwig files were created with deepTools version 3.4.2 (64) bamCoverage tool with the parameters -bs 1 -p max -normalizeUsing RPKM.

### ChIP-seq analysis

K562 Cyclin T1 and RNAPII ChIP-seq spiked with mouse cells were obtained from GSE148433 (30). HeLa SF3B1 MNase-seq and its Input were obtained from GSE65644 (54). Adapters were trimmed with Cutadapt in paired-end mode with the options: --minimum-length 10 -q 15, 10-j 16 -A GATCGTCGGACTGTAGAACTCTGAAC -a AGATCGGAAGAGCACACGTCTGAACTCCAGTCA. Trimmed reads were mapped to the human GRCh38.p13 and to the mouse GRCm38 reference genomes with STAR and the parameters: --runThreadN 16 --readFilesCommand gunzip -c -k -alignIntronMax 1 -- outFilterMultimapNmax 1 --limitBAMsortRAM 20000000000 --outSAMtype BAM SortedByCoordinate. SAMtools was used to keep properly paired and mapped reads and to remove PCR duplicates. Reads mapping to the DAC Exclusion List Regions (accession: ENCSR636HFF) (65) were removed with BEDtools version 2.29.2 (66). For HeLa SF3B1 MNase-seq, FPKM-normalised bigwig files were created with deepTools bamCoverage tool with the parameters -bs 10 -e 150 -p max -normalizeUsing RPKM. For K562 Cyclin T1 and RNAPII ChIP-seq, SAMtools view with the -s option was used to subsample all the bam files to the bam file containing the lowest number of reads. RNAPII PlaB replicate 2 and its associated Input was removed at this step due to a low number of mapped reads. The normalisation factor was then calculated as: (number of mouse reads) / (number of mouse + number of human reads) and applied to the generation of the bigwig files with deepTools bamCoverage tool with the parameters -bs 10 -p max -e -scaleFactor 1 / (normalisation factor).

### Quantification and RNAPII pausing index

Quantification of RNAPII and Cyclin T1 ChIP-seq signal over promoter regions (defined as TSS – 250 bp to TSS + 250 bp) and gene body (defined as TSS + 500 bp to poly(A) site) was performed with bedtools multicov and normalised with the spike-in normalisation factor defined in “ChIP-seq analysis”. The pausing index is defined as: (Quantification “Promoter region” / 500) / (Quantification “Gene body” / Size_(Gene body)_). A PlaB / DMSO ratio ≤ 0.5 is considered decreased, ≥ 2 increased, and between 0.5 and 2 unchanged.

### □ Metagene and boxplots

Metagene profiles were generated with Deeptools2 computeMatrix tool with the parameters-bs 10 -p max -m 4000 (for scaled regions) and–maxThreshold 2500 parameter added for mNET-seq analysis. The plotting data were obtained with plotProfile –outFileNameData. Metagene profiles representing the (IP / Input) signal (ChIP□seq) or the mNET□seq signal and boxplots (min to max, with the box representing first, median, and third quartile) were plotted with GraphPad Prism 9.5.

### Statistical tests

Statistical tests are indicated in figures legends and were performed with GraphPad Prism 9.5.

## RESULTS

### Rapid inhibition of SF3B1 decreases pre-mRNA splicing and RNAPII transcription

To investigate the effect of rapid inhibition of the U2 snRNP factor SF3B1 by PlaB (29) or HB (55), we first carried out RT-PCR analysis of splicing of BRD2 exons 4 and 5 and *DNAJB1* exons 2 and 3 over a time-course of 30 min to 4 h treatment of HeLa and HEK293 cells with 1 µM PlaB or 1 µM HB (Figure 1A and Supplementary Figure 1A and B). As previously observed (30), we found that after 30 min, SF3B1 inhibition increases intron retention. Concomitant to the loss of splicing, SF3B1 inhibition with PlaB decreases 5’EU incorporation in nascent RNA and RNAPII CTD Ser2 phosphorylation, a mark of transcription elongation (5), albeit not as strongly as CDK9 inhibition with the CDK9 inhibitor DRB (Figure 1B-E and Supplementary Figure 1C-H). These results confirm that a short inhibition of SF3B1 with PlaB is sufficient to affect both splicing and RNAPII transcription.

**Figure 1.**
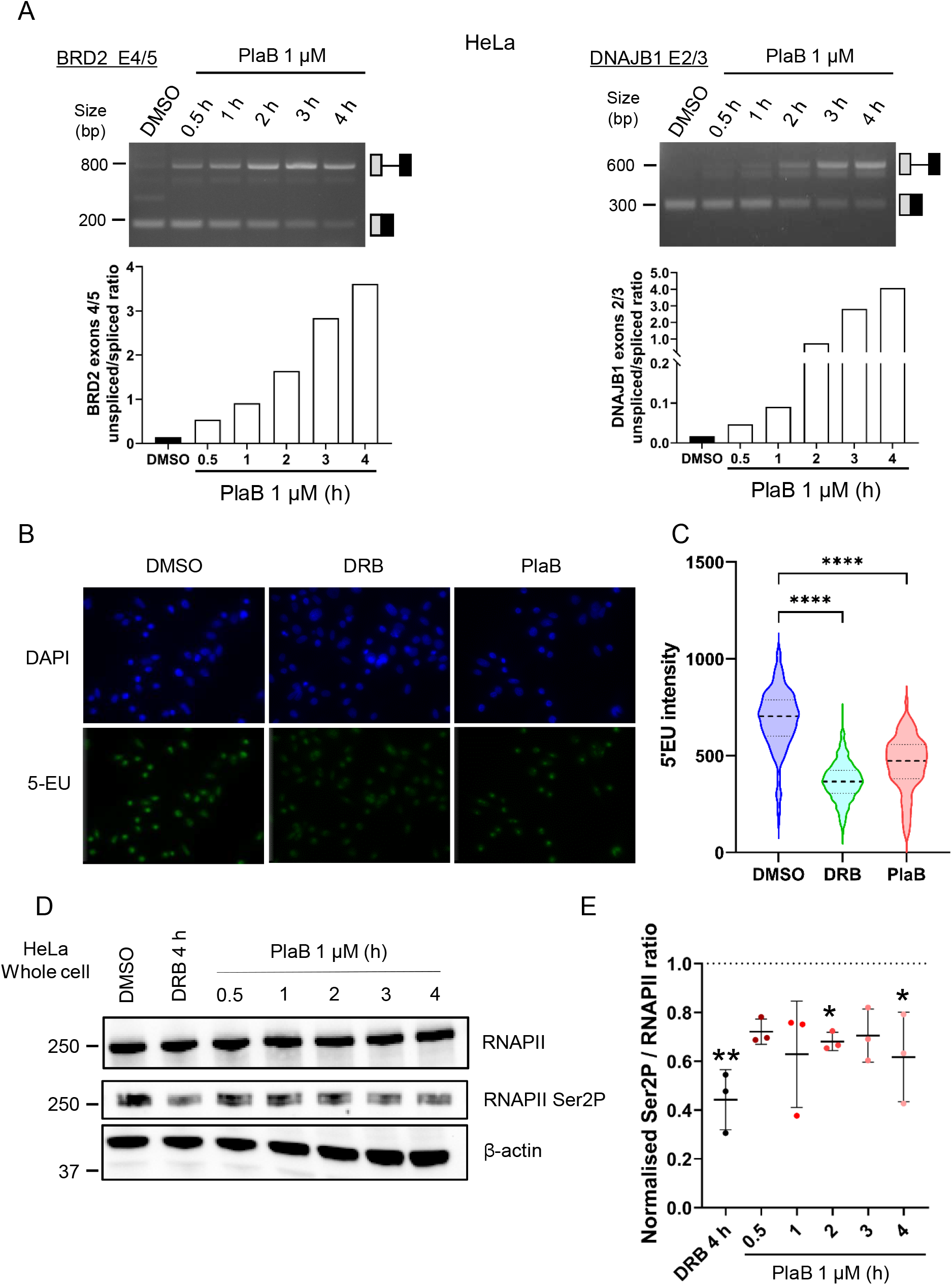
Rapid inhibition of SF3B1 decreases pre-mRNA splicing and RNAPII transcription. (**A**) RT-PCR with primers amplifying the intron located between exons 4 and 5 of BRD2 or exons 2 and 3 of DNAJB1. HeLa cells were treated with DMSO or 1 µM PlaB for 30 min to 4 h. The location of the spliced and unspliced RNA is shown on the left of the panel. The percentages of unspliced RNA compared to total (spliced + unspliced) are shown below. (**B**) Representative images of immunofluorescence analysis of 5’EU incorporation in HeLa cells treated with DMSO, 100 µM DRB, a CDK9 inhibitor, or 1 µM PlaB for 1h. EU (green), DAPI (blue), scale bars: 50 µm. (**C**) Quantification of 5’EU intensity per nucleus for DMSO (blue), DRB (green), and PlaB (red). Boxplot settings are: min to max values with the box showing 25-75 percentile range. 394 nuclei were quantified per condition. Statistical test: Kruskal-Wallis test. P-value: **** < 0.0001. (**D**) Representative whole cell western blots of total RNAPII, RNAPII Ser2 phosphorylation, and β-actin (loading control) from HeLa cells treated with DMSO, 100 µM DRB for 4 h, or 1 µM PlaB for 30 min to 4h. (**E**) Quantification of the western blots shown in (D) presented as RNAPII Ser2P / total RNAPII following normalisation to the loading control and to the DMSO condition. Statistical test: Kruskal-Wallis test, n = 3 biological replicates. P-value: * < 0.05, ** < 0.01.

### SF3B1 inhibition increases RNAPII pausing

To understand how rapid inhibition of SF3B1 decreases 5’EU incorporation and RNAPII CTD Ser2 phosphorylation, we performed total, Ser2P, and Ser5P RNAPII mNET-seq (32) in HeLa cells treated with DMSO or 1 µM PlaB for 30 min (Figure 2A-C and Supplementary Figure 2A and B). Consistent with published results (30,34), we observed an increase in RNAPII pausing and a decrease in transcription elongation following PlaB treatment. Analysis of the Ser5P mNET-seq signal on the last nucleotide of internal exons, which is associated with co-transcriptional cleavage of the RNA due to pre-mRNA splicing (32), shows, as previously described (67), a decrease following PlaB treatment, indicative of reduced RNA cleavage during co-transcriptional splicing (Supplementary Figure 2C and D). The effect of SF3B1 inhibition on RNAPII transcription was confirmed by RNAPII ChIP-qPCR on the model gene *KPNB1* following different treatment times with PlaB and HB (Figure 2D). We also observed that in addition to the effect on intron-containing genes, PlaB treatment causes an increase in RNAPII pausing on intronless genes, an observation previously made using spiked total RNAPII ChIP-seq in K562 cells treated with PlaB for 1 h (Supplementary Figure 2E and F) (30).

**Figure 2.**
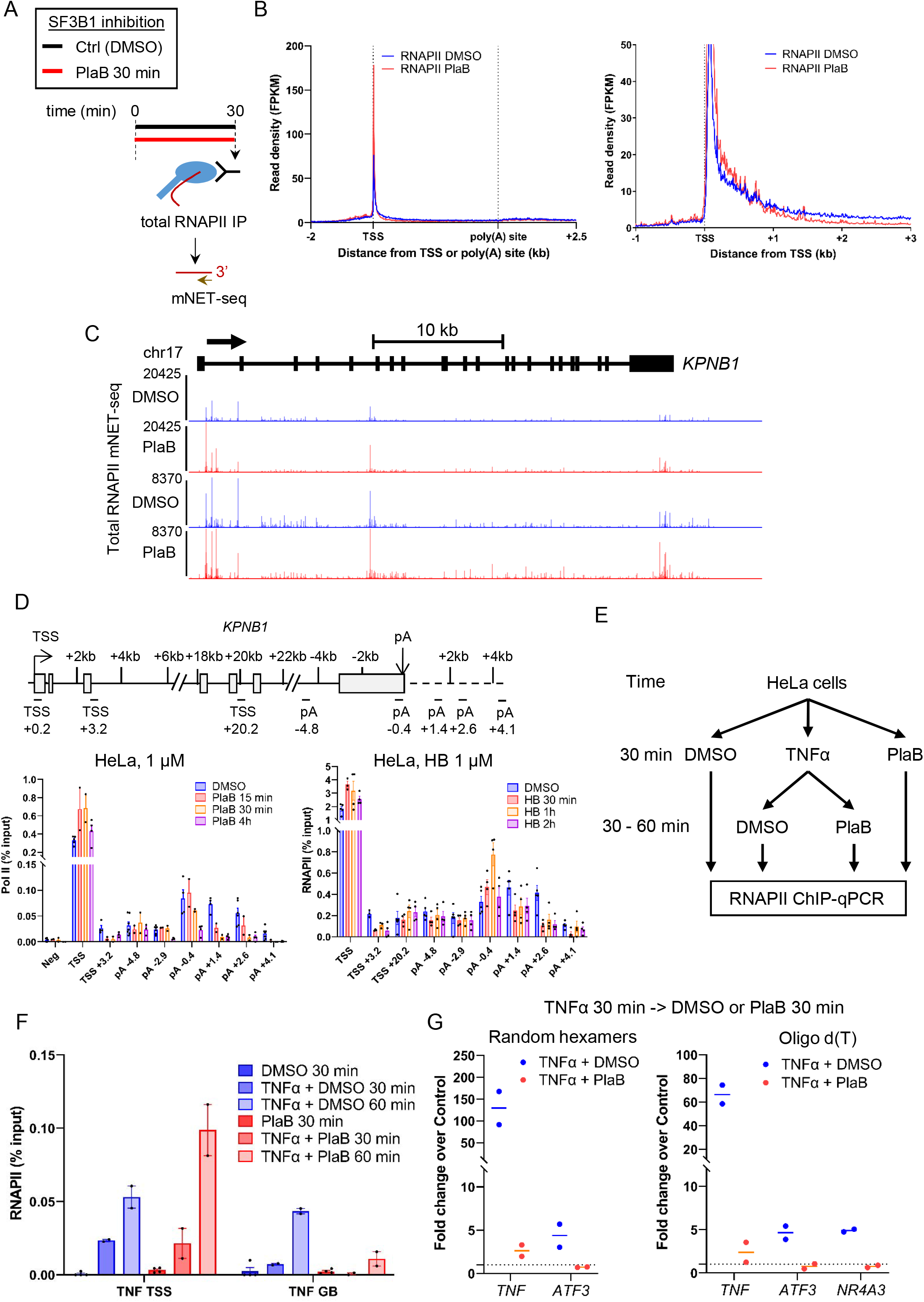
SF3B1 inhibition increases RNAPII pausing. (**A**) Schematic of the mNET-seq technique. (**B**) Metagene profile of total RNAPII mNET-seq in HeLa cells treated with DMSO (blue) or 1 µM PlaB (red) for 30 min on scaled expressed protein-coding genes (left) and on TSS -1 kb to TSS + 3 kb of expressed protein-coding genes (right). (**C**) Screenshot of the genome browser for total RNAPII mNET-seq DMSO (blue) and PlaB (red) tracks of the protein-coding gene *KPNB1*. The arrow indicates the sense of transcription. (**D**) Total RNAPII ChIP-qPCR across the *KPNB1* gene in HeLa cells treated with DMSO, 1 µM PlaB for 15 min, 30 min, or 4 h, or with 1 µM HB for 30 min, 1h, or 2h. Statistical test: Unpaired t-test, n = 4 biological replicates. P-value: * < 0.05, ** < 0.01. (**E**) Schematic of the TNFα induction experiments in HeLa cells. (**F**) Total RNAPII ChIP-qPCR on *TNF* TSS and gene body gene in HeLa cells treated with DMSO, 100 ng/ml TNFα, and/or 1 µM PlaB, n = 2 biological replicates. (**G**) qRT-PCR performed with random hexamers or oligo d(T) of TNF, ATF3, and NR4A3 in HeLa cells treated with DMSO, 100 ng/ml TNFα, and/or 1 µM PlaB, n = 2 biological replicates.

To investigate whether RNAPII pause release is affected by SF3B1 inhibition, we analysed the effect of PlaB on transcription of TNFα-induced genes. HeLa cells were treated for 30 min with DMSO, 10 ng/ml TNFα, or 1 µM PlaB and then the TNFα-treated cells were treated for a further 30 or 60 min with DMSO or 1 µM PlaB before performing total RNAPII ChIP-qPCR on the promoter and gene body of the TNF gene, which is induced by TNFα (Figure 2E and F).

Interestingly, TNFα treatment followed by DMSO treatment induces transcription initiation and elongation whereas TNFα treatment followed by PlaB treatment promotes transcription initiation and RNAPII pausing but poor transcription elongation, indicating a requirement for SF3B1 activity to promote RNAPII pause release. These results were confirmed by random hexamers and oligo(dT) qRT-PCR of the TNFα-inducible genes *TNF, ATF3*, and *NR4A3*, which indicates that transcription is induced in the TNFα 30 min + DMSO 30 min but poorly after TNFα 30 min + PlaB 30 min treatment (Figure 2G). To determine whether the effect of SF3B1 inhibition on transcriptional activity is due to cellular stress, as previously observed for U2AF1 S34F and Q157R mutants (68), we analysed the level of G3BP1, a marker of cellular stress granules, by immunofluorescence, (69), following 30 min treatment with DMSO, PlaB, and HB. There was no increase in this marker (Supplementary Figure 2G and H), indicating that the effect of SF3B1 inhibition on RNAPII transcription, at least after 30 min treatment, is not likely to be due to cellular stress caused by splicing inhibition.

### SF3B1 and CDK9 inhibition decrease transcription downstream of histone gene bodies

As RNAPII transcription and Cyclin T1 recruitment are affected by PlaB treatment on protein-coding genes, including intronless genes, we also investigated the effect of SF3B1 inhibition on non-intron-containing replication-activated histone genes by also re-analysing additional published datasets (Supplementary Figure 3A-D). We observed that while Cyclin T1 recruitment is decreased by PlaB treatment, RNAPII mNET-seq, ChIP-seq, and ChIP-qPCR do not show any major effect on transcription across histone gene bodies after SF3B1 inhibition. However, we found that both SF3B1 and CDK9 inhibition, with either a Raji CDK9 analogue-sensitive cell line (60) or with the inhibitor DRB in HeLa cells (38), decrease transcription downstream of histone gene bodies (Supplementary Figure 3E-H), reminiscent of the premature transcription termination induced by CDK9 inhibition on other protein-coding genes (38,70). These results indicate that SF3B1 inhibition affects transcription of all protein-coding genes, not only intron-containing genes.

**Figure 3.**
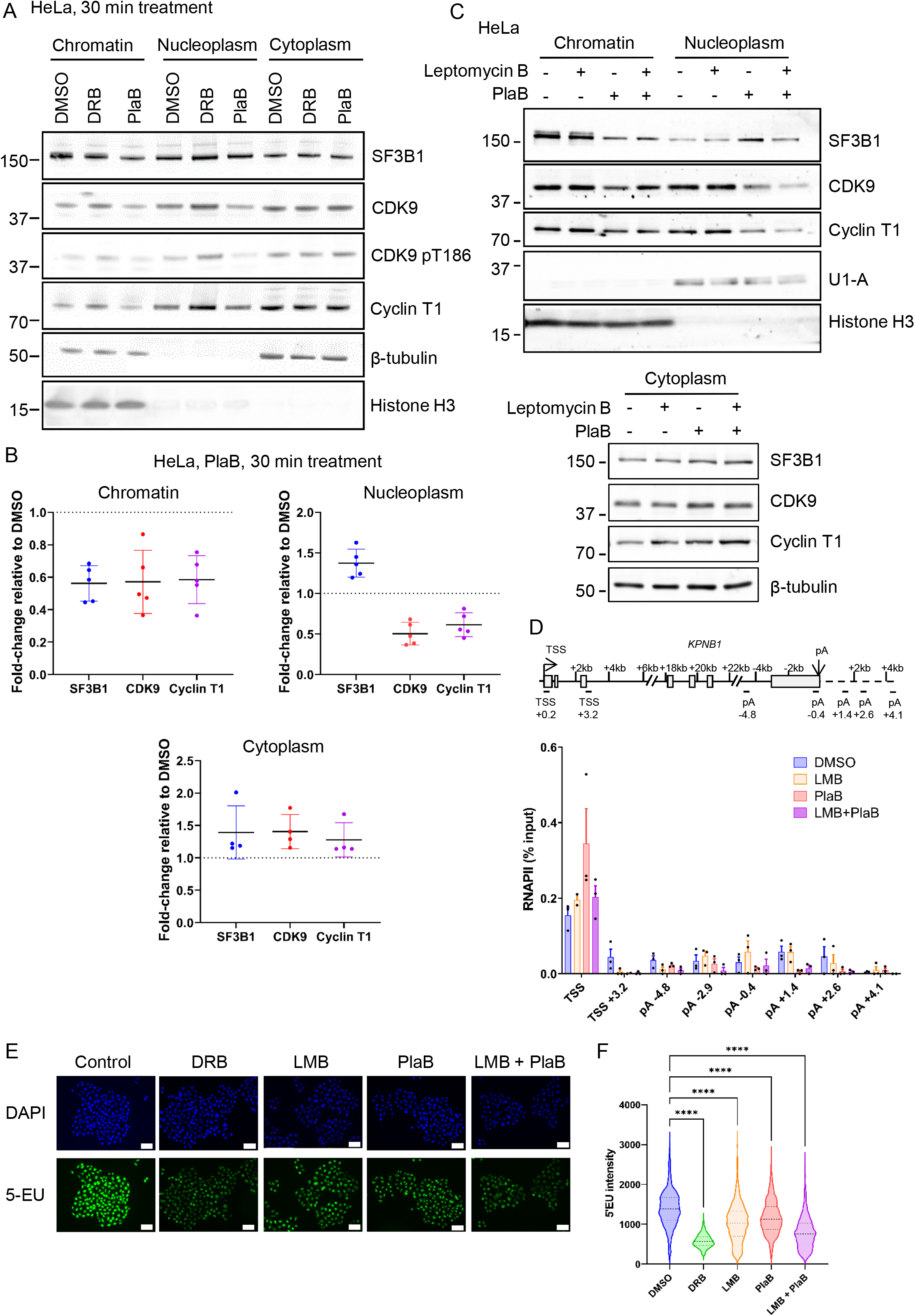
SF3B1 inhibition increases P-TEFb nuclear export. (**A**) Representative western blots of SF3B1, CDK9, CDK9 T186 phosphorylation, Cyclin T1, β-tubulin (loading control), and histone H3 (loading control) from chromatin, nucleoplasm, and cytoplasm fractions of HeLa cells treated with DMSO, 100 µM DRB, and 1 µM PlaB for 30 min. (**B**) Quantification of western blots of SF3B1, CDK9, and Cyclin T1 from chromatin, nucleoplasm, and cytoplasm fractions of HeLa cells following treatment with 1 µM PlaB for 30 min. The quantifications have been normalised to loading control and to DMSO, n = 4-5 biological replicates (including PlaB 30 min from Supplementary Figure 5C and PlaB from Supplementary Figure 6D). (**C**) Representative western blots of SF3B1, CDK9, Cyclin T1, U1-A (loading control), β-tubulin (loading control), and histone H3 (loading control) from chromatin, nucleoplasm, and cytoplasm fractions of HeLa cells treated with DMSO, ethanol, 0.02 µM Leptomycin B, or 1 µM PlaB. Treatments were performed as follow: Ethanol 1h/DMSO 30 min (-/-), LMB 1h/DMSO 30 min (+/-), Ethanol 1h/PlaB 30 min (-/+), LMB 1h/PlaB 30 min (+/+). (**D**) Total RNAPII ChIP-qPCR across the *KPNB1* gene in HeLa cells treated with DMSO, ethanol, 0.02 µM Leptomycin B, or 1 µM PlaB as described in (C), n = 3 biological replicates. (**E**) Representative images of immunofluorescence analysis of 5’EU incorporation in HeLa cells treated with DMSO, ethanol, 100 µM DRB, 0.02 µM Leptomycin B, or 1 µM PlaB as described in (C). EU (green), DAPI (blue), scale bars: 50 µm. (**F**) Quantification of 5’EU intensity per nucleus for DMSO (blue), DRB (green), LMB (orange), PlaB (red), and LMB & PlaB (purple). Boxplot settings are: min to max values with the box showing 25-75 percentile range. 1,620 nuclei were quantified per condition. Statistical test: Kruskal-Wallis test. P-value: **** < 0.0001.

### SF3B1 inhibition increases P-TEFb nuclear export

PlaB treatment clearly causes a reduction of P-TEFb close to the TSS of protein-coding genes (30). To determine the fate of P-TEFb after SF3B1 inhibition, we performed chromatin, nucleoplasm, and cytoplasm cellular fractionation followed by western blotting of SF3B1, CDK9, and Cyclin T1 after treatment with DMSO, 100 µM DRB, or 1 µM PlaB for 30 min (Figure 3A and B). Contrary to CDK9 inhibition with DRB and other CDK9 inhibitors, which are known to increase P-TEFb association with chromatin (71,72), PlaB treatment results in a partial loss of P-TEFb from the chromatin and nucleoplasm fractions and an associated increase in the cytoplasm (Figure 3A and B). PlaB also causes SF3B1 to move from chromatin to nucleoplasm and cytoplasm. These results were confirmed with a PlaB treatment time course from 30 min to 4 h, with the strongest loss of P-TEFb chromatin association occurring after 30 min of treatment (Supplementary Figure 4A-C). The nuclear CDK9 signal also significantly decreases as measured by immunofluorescence following treatment with 1 µM PlaB for 30 min (Supplementary Figure 4D and E).

**Figure 4.**
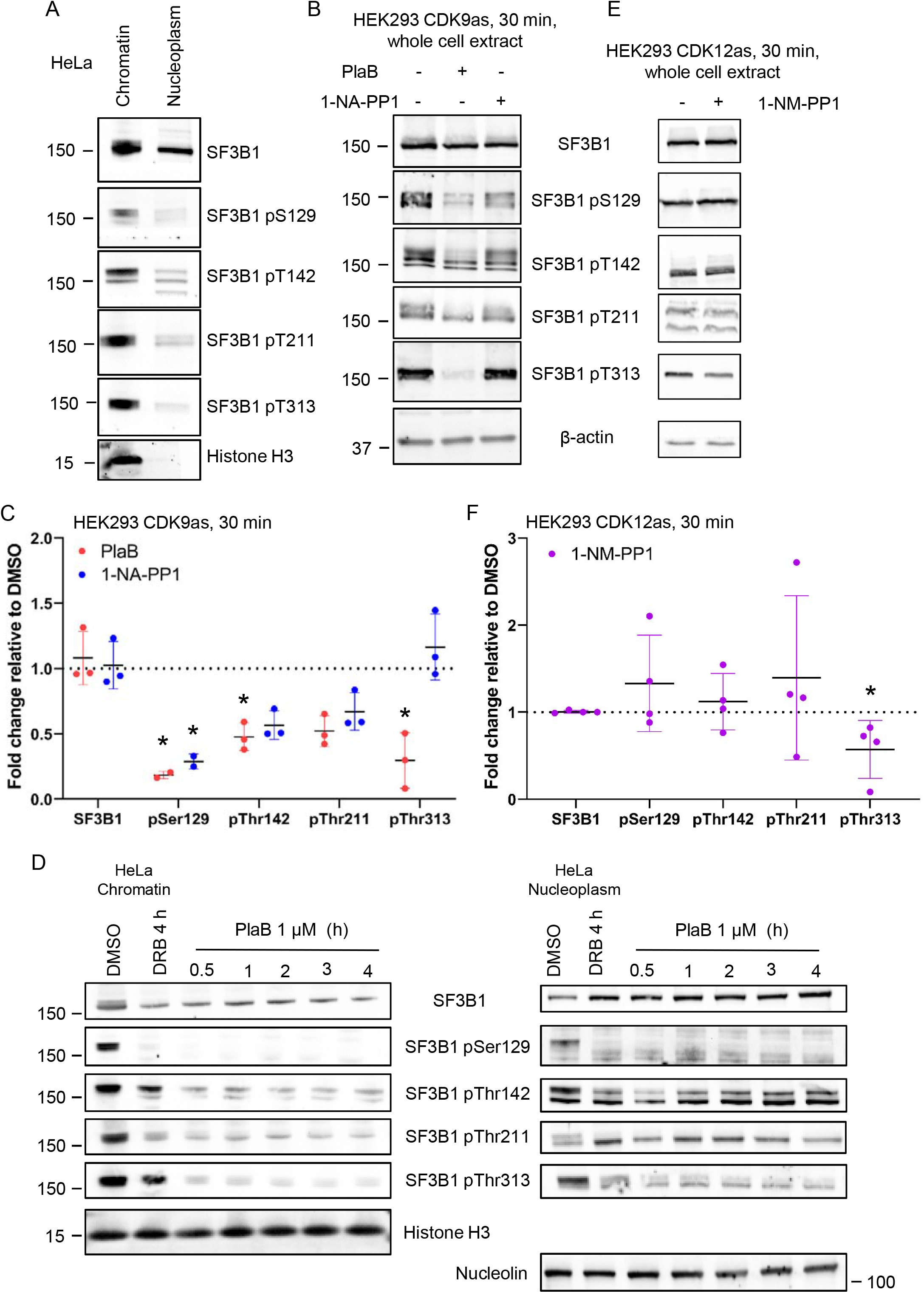
SF3B1 and CDK9 inhibition decrease phosphorylation and chromatin association of SF3B1. (**A**) Western blots of total SF3B1, SF3B1 S129, T142, T211, and T313 phosphorylation, and histone H3 (loading control) from chromatin and nucleoplasm fractions of untreated HeLa cells. (**B**) Representative western blots of total SF3B1, SF3B1 S129, T142, T211, and T313 phosphorylation, and β-actin (loading control) from whole cell extract of HEK293 CDK9 analogue sensitive cells treated with DMSO, 1 µM PlaB or 15 µM 1-NA-PP1 for 30 min. (**C**) Quantification of western blots of total SF3B1 and SF3B1 S129, T142, T211, and T313 phosphorylation from whole cell extract of HEK293 CDK9 analogue sensitive cells treated with 1 µM PlaB or 15 µM 1-NA-PP1 for 30 min. The quantifications have been normalised to loading control and to DMSO. Statistical test: Kruskal-Wallis test, n = 2 - 3 biological replicates. P-value: * < 0.05. (**D**) Western blots of total SF3B1, SF3B1 S129, T142, T211, and T313 phosphorylation, histone H3 (loading control), and nucleolin (loading control) from chromatin and nucleoplasm fractions of HeLa cells treated with DMSO, 100 µM DRB for 4 h, or with 1 µM PlaB for 30 min to 4h. (**E**) Western blots of total SF3B1, SF3B1 S129, T142, T211, and T313 phosphorylation, and β-actin (loading control) from whole cell extract of HEK293 CDK12 analogue sensitive cells treated with DMSO or 7.5 µM 1-NM-PP1 for 30 min. Two biological replicates shown. (**F**) Quantification of western blots of total SF3B1 and SF3B1 S129, T142, T211, and T313 phosphorylation from whole cell extract of HEK293 CDK12 analogue sensitive cells treated with 7.5 µM 1-NM-PP1 for 30 min. The quantifications have been normalised to loading control and to DMSO. Statistical test: Kruskal-Wallis test, n = 4 biological replicates. P-value: * < 0.05.

To test whether SF3B1 inhibition causes degradation of SF3B1, CDK9, or Cyclin T1, we performed whole cell extract western blotting across a PlaB treatment time course of 30 min to 4 h (Supplementary Figure 5A and B). The global level of these factors remains stable over the time course. In addition, western blotting on chromatin and nucleoplasmic fractions performed in the absence or presence of 1 µM PlaB and/or 1 µM Bortezomib, a nuclear proteasome inhibitor (73), confirmed that SF3B1 inhibition does not promote degradation of SF3B1, CDK9, and Cyclin T1 (Supplementary Figure 5C).

**Figure 5.**
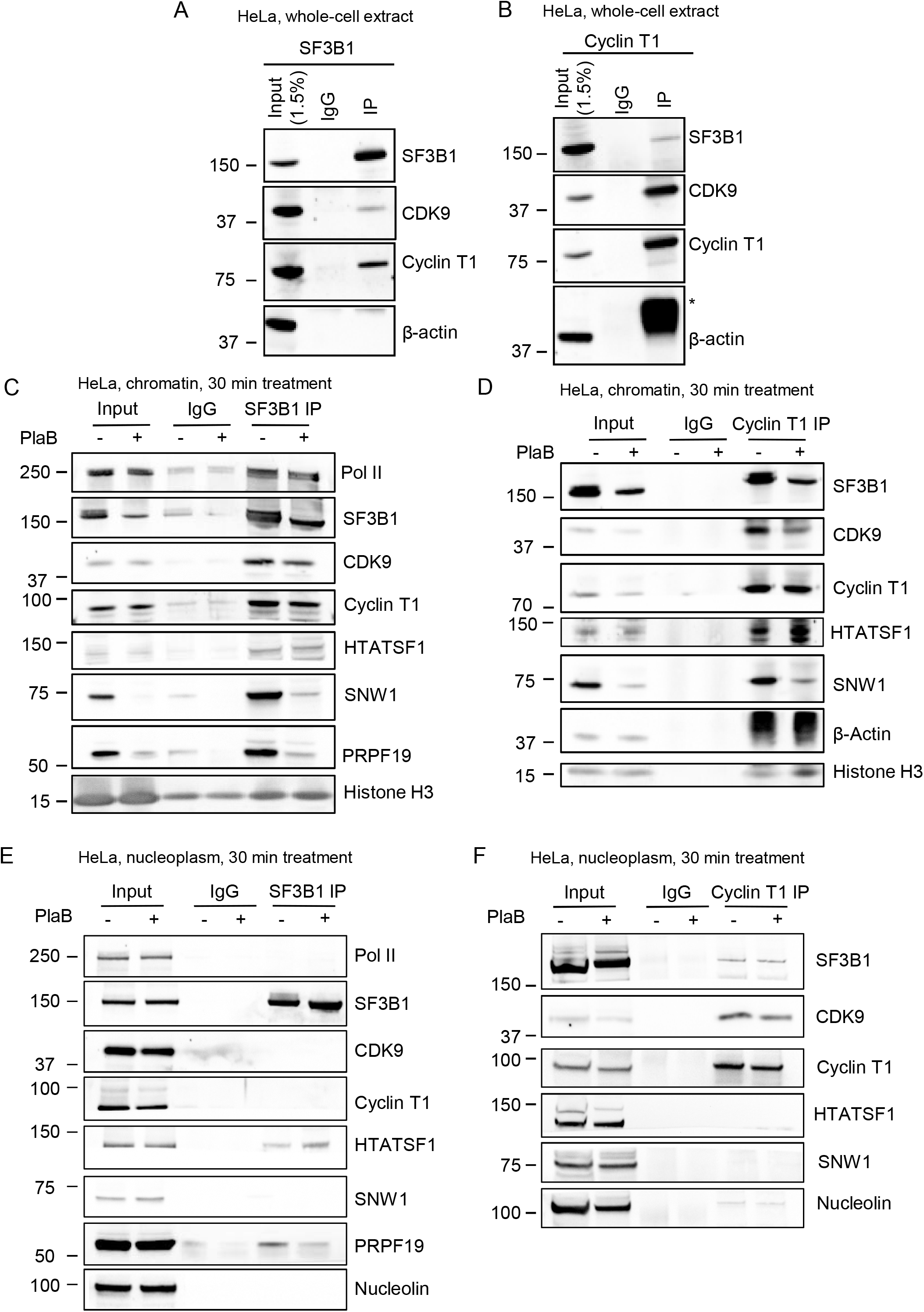
SF3B1 inhibition affects predominantly SNW1 in the chromatin-associated SF3B1/HTATSF1/SNW1/P-TEFb complex. (**A**) Co-immunoprecipitation of SF3B1 from whole cell extract of untreated HeLa cells followed by western blots of SF3B1, CDK9, Cyclin T1, and β-actin (negative control). (**B**)Co-immunoprecipitation of Cyclin T1 from whole cell extract of untreated HeLa cells followed by western blots of SF3B1, CDK9, Cyclin T1, and β-actin (negative control). *: IgG. (**C**) Co-immunoprecipitation of SF3B1 from chromatin of HeLa cells treated with DMSO or 1 µM PlaB for 30 min followed by western blots of total RNAPII, SF3B1, CDK9, Cyclin T1, HTATSF1, SNW1, PRPF19, and histone H3. (**D**) Co-immunoprecipitation of Cyclin T1 from chromatin of HeLa cells treated with DMSO or 1 µM PlaB for 30 min followed by western blots of SF3B1, CDK9, Cyclin T1, HTATSF1, SNW1, β-actin, and histone H3. (**E**) Co-immunoprecipitation of SF3B1 from nucleoplasm of HeLa cells treated with DMSO or 1 µM PlaB for 30 min followed by western blots of total RNAPII, SF3B1, CDK9, Cyclin T1, HTATSF1, SNW1, PRPF19, and Nucleolin. (**F**) Co-immunoprecipitation of Cyclin T1 from nucleoplasm of HeLa cells treated with DMSO or 1 µM PlaB for 30 min followed by western blots of SF3B1, CDK9, Cyclin T1, HTATSF1, SNW1, and Nucleolin.

To determine whether we could block P-TEFb nuclear export, we treated HeLa cells with PlaB and/or Leptomycin B (LMB), an inhibitor of the nuclear export protein CRM1 (74), and performed western blotting of SF3B1, CDK9, and Cyclin T1 on chromatin, nucleoplasmic, and cytoplasmic fractions (Figure 3C and Supplementary Figure 5D). We found that a one-hour pre-treatment with LMB is not sufficient to block P-TEFb nuclear export due to the mechanism of action of LMB (see Discussion). However, we observed that P-TEFb loss from chromatin following PlaB treatment is reduced in presence of LMB. We therefore wondered whether this reduced loss of P-TEFb caused by LMB treatment abrogates the effect of PlaB on RNAPII transcription. To test this, we performed RNAPII ChIP-qPCR and qRT-PCR on our model ∼40 kb gene *KPNB1* (Figure 3D and Supplementary Figure 5E). While PlaB treatment causes the expected RNAPII profile and loss of nascent transcripts, a one-hour treatment with LMB reduces RNAPII entering productive elongation (TSS +3.2 kb primer) without affecting RNAPII pausing and termination. The combination of PlaB and LMB has a limited reduced effect on RNAPII pausing but a similar decrease in RNAPII transcription across the gene body as PlaB alone. We confirmed these results with 5’EU incorporation experiments of cells treated with DMSO, 100 µM DRB, 1 µM LMB, 1 µM PlaB, and 1 µM LMB and 1 µM PlaB (Figure 3E and F). LMB treatment by itself reduces 5’EU incorporation and the combination of LMB and PlaB further decreases 5’EU incorporation compared to LMB or PlaB alone, indicating a potential additive effect. As the 5’EU signal also includes RNAPI rDNA transcription, we also tested pre-rRNA transcript levels and found that LMB, but not PlaB or LMB & PlaB, reduces rDNA transcription (Supplementary Figure 5F).

These results indicate that SF3B1 inhibition promotes a rapid increase in P-TEFb nuclear export and a reduction in SF3B1 binding to chromatin. Blocking nuclear export of P-TEFb by LMB does not reverse the effect on transcription.

### Inhibition of SF3B1 or CDK9 decreases phosphorylation and chromatin association of SF3B1

While SF3B1 inhibition results in the loss of SF3B1 and P-TEFb from chromatin, SF3B1 phosphorylation also regulates SF3B1 chromatin association (39,52) and P-TEFb phosphorylates SF3B1 (37,38). To understand better the links between SF3B1 chromatin association, SF3B1 phosphorylation, and P-TEFb, we first investigated the cellular localisation of different phosphoforms of SF3B1 by analysing phosphorylation of SF3B1 residues Ser129, Thr142, Thr211, and Thr313 by western blotting of chromatin and nucleoplasm fractions (Figure 5A). In agreement with previous findings, we observed that phosphorylation of these SF3B1 residues is highest on chromatin (39,52). To determine which of the four SF3B1 phosphorylated residues is affected by CDK9 and SF3B1 inhibition, we used our previously published HEK293 CDK9 analogue sensitive cell line (38) and treated the cells for 30 min with either 15 µM 1-NA-PP1 or 1 µM PlaB (Figure 5B and C). Whole-cell western blotting indicates that SF3B1 or CDK9 inhibition does not affect the level of SF3B1 protein but causes a decrease in phosphorylation of Ser129, Thr142, Thr211, and in the case of PlaB, also of Thr313. To test whether SF3B1 present on chromatin after PlaB treatment is still phosphorylated on these four residues, we performed western blotting of SF3B1 and the different phospho-residues on HeLa chromatin and nucleoplasm fractions following CDK9 inhibition with 100 µM DRB for 4 h or with 1 µM PlaB across a time course of 30 min to 4 h (Figure 5D). We found that following PlaB treatment, the SF3B1 still associated with chromatin has lost phosphorylation of these four residues. The loss of phosphorylated SF3B1 is also noticeable by the disappearance of the top SF3B1 band visible in the DMSO condition on chromatin, which likely corresponds to hyperphosphorylated SF3B1.

Another transcriptional kinase, CDK12, was also found to phosphorylate SF3B1 when inhibited with the CDK12/CDK13 inhibitor THZ531 (43). Using our CDK12 analogue sensitive cell line (56), we observed that inhibition of CDK12 decreases phosphorylation of Thr313, but not of Ser129, Thr142, or Thr211 (Figure 4E and F). These results show an interdependent regulation between P-TEFb and SF3B1 and further emphasize the intimate relationship between SF3B1 and transcriptional CDKs.

### SF3B1 inhibition affects predominantly SNW1 in the chromatin-associated SF3B1/HTATSF1/SNW1/P-TEFb complex

As PlaB treatment reduces SF3B1 and P-TEFb on chromatin, we investigated whether SF3B1, CDK9, and Cyclin T1 interact. We first preformed HeLa whole cell SF3B1, CDK9, and Cyclin T1 co-immunoprecipitation experiments followed by western blotting (Figure 5A and B and Supplementary Figure 6A). In agreement with (75) but contrary to (42), we found interaction between SF3B1 and P-TEFb. To understand better in which cellular fraction, whether it is mediated via HTATSF1 and/or SNW1, and the effect of PlaB on these interactions, we performed SF3B1 and Cyclin T1 co-immunoprecipitation from HeLa chromatin (Figure 5C and D), nucleoplasmic (Figure 5E and F), and cytoplasmic (Supplementary Figure 6B and C) fractions treated for 30 min with DMSO or PlaB. We found that the interaction between SF3B1 and P-TEFb occurs on chromatin but not in the nucleoplasm and cytoplasm. Additionally, we found that SF3B1 and P-TEFb interacts with HTATSF1 and SNW1 on chromatin but that in the nucleoplasm and cytoplasm, only an interaction between SF3B1 and HTATSF1 is detected. Following PlaB treatment, we observed, as expected, that interaction between SF3B1 and PRPF19, a member of the NineTeen Complex (76), which is part of the spliceosome B complex, is lost following PlaB treatment, which blocks the spliceosome in the A-like complex (77). Interestingly, the interactions between SF3B1, HTATSF1, CDK9, and Cyclin T1 still occur on chromatin after PlaB treatment, albeit with some reduction in SF3B1 and P-TEFb association. The interaction between SF3B1 and HTATSF1 in the nucleoplasm and cytoplasm is not affected by SF3B1 inhibition. Conversely, SF3B1 or Cyclin T1 interaction with SNW1 is lost on chromatin following PlaB treatment, likely due to the reduction of SNW1 on chromatin following SF3B1 inhibition (see Input, Figure 5C and D). Immunoprecipitation of HTATSF1 from whole HeLa cells treated for 30 min with DMSO or PlaB followed by western blotting confirmed the results obtained with SF3B1 and Cyclin T1 co-immunoprecipitation (Supplementary Figure 6D).

**Figure 6.**
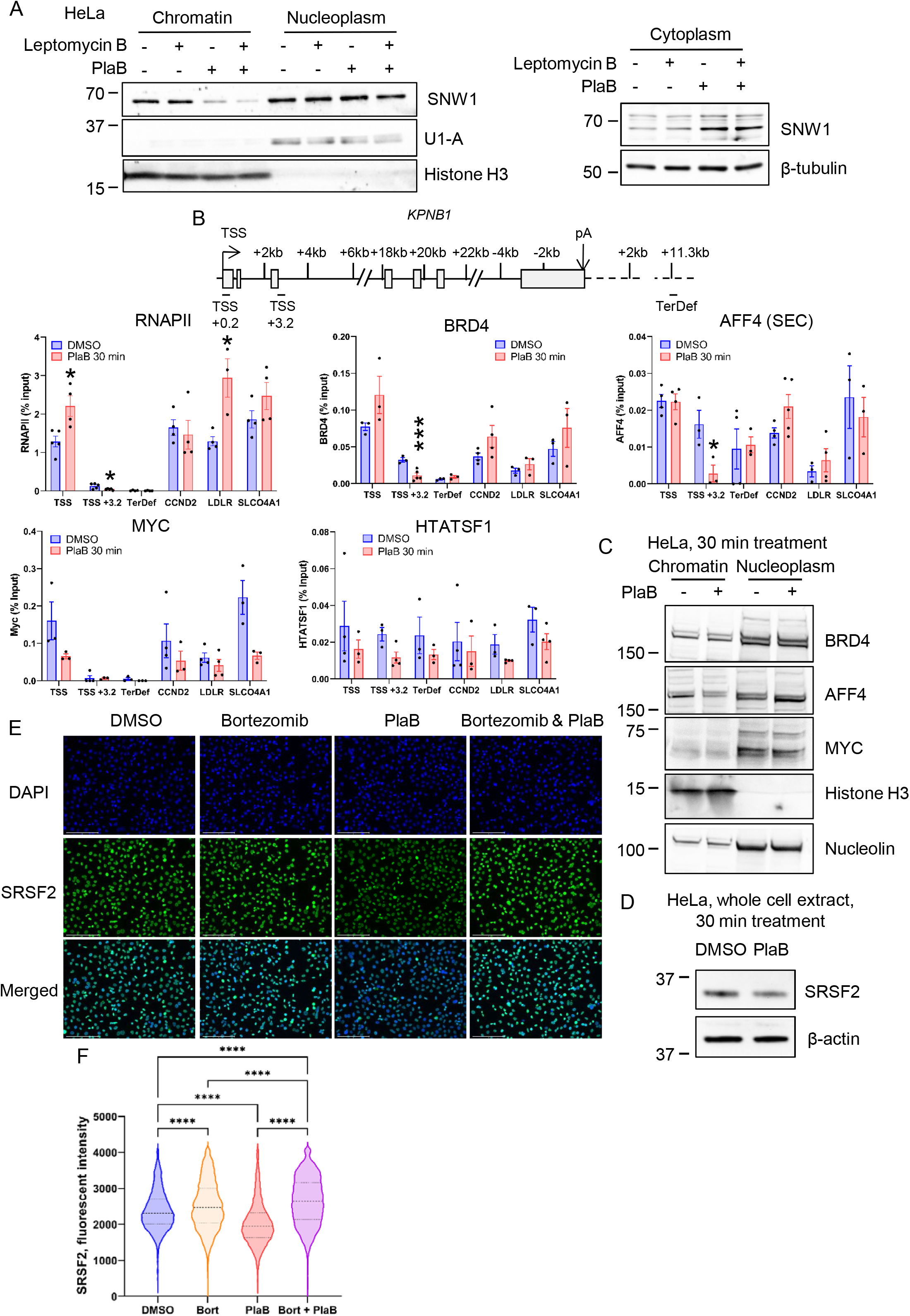
SF3B1 inhibition promotes SNW1 nuclear export, AFF4 loss from chromatin, and SRSF2 degradation. (**A**)Western blots of SNW1, U1-A (loading control), histone H3 (loading control), and β-tubulin (loading control) from chromatin, nucleoplasm, and cytoplasm fractions of HeLa cells treated with DMSO, ethanol, 0.02 µM Leptomycin B, or 1 µM PlaB. Treatments were performed as follow: Ethanol 1h/DMSO 30 min (-/-), LMB 1h/DMSO 30 min (+/-), Ethanol 1h/PlaB 30 min (-/+), LMB 1h/PlaB 30 min (+/+). (**B**) Total RNAPII, BRD4, AFF4, MYC, and HTATSF1 ChIP-qPCR on the promoter regions of *KPNB1, CCND2, LDLR*, and *SLCO4A1* in HeLa cells treated with DMSO or 1 µM PlaB for 30 min. Statistical test: Unpaired t-test, n = 3-4 biological replicates. P-value: * < 0.05, *** < 0.001. The RNAPII ChIP-qPCR data for TSS, TSS+3.2, LDLR, and SLCO4A1 are the same as the ones shown in Supplementary Figure 7C (same ChIP-qPCR experiments). (**C**) Western blots of NRD4, AFF4, MYC, histone H3 (loading control), and Nucleolin (loading control) from chromatin and nucleoplasm fractions of HeLa cells treated with DMSO or 1 µM PlaB for 30 min. (**D**) Western blots of SRSF2 and β-actin (loading control) from whole cell extract of HeLa cells treated with DMSO or 1 µM PlaB for 30 min. (**E**) Representative images of immunofluorescence analysis of SRSF2 in HeLa cells treated with DMSO, 1 µM Bortezomib, or 1 µM PlaB. Treatments were performed as follow: DMSO 1h/DMSO 30 min (DMSO), Bortezomib 1h/DMSO 30 min (Bortezomib), DMSO 1h/PlaB 30 min (PlaB), Bortezomib 1h/PlaB 30 min (Bortezomib & PlaB). SRSF2 (green), DAPI (blue), scale bars: 50 µm. (**F**) Quantification of SRSF2 fluorescent intensity for DMSO (blue), Bortezomib (orange), PlaB (red), and Bortezomib & PlaB (purple). Boxplot settings are: min to max values with the box showing 25-75 percentile range. 10,106 nuclei were quantified per condition. Statistical test: Kruskal-Wallis test. P-value: **** < 0.0001.

These results indicate that P-TEFb interacts with SF3B1, HTATSF1, and SNW1 on chromatin and that SNW1, and to a lesser extent P-TEFb, are lost from the complex after PlaB treatment.

### SF3B1 inhibition promotes SNW1 nuclear export, AFF4 loss from chromatin, and SRSF2 degradation

As we found that PlaB treatment reduces the association of SNW1 with chromatin, we investigated whether SNW1 is exported to the cytoplasm by performing western blotting on different cellular fractions (Figure 6A). Comparable to P-TEFb, we found that SF3B1 inhibition results in a clear loss of SNW1 from chromatin, but not from the nucleoplasm, and in an accumulation of SNW1 in the cytoplasm, which is not blocked by a 1h pre-treatment with LMB.

To determine whether other P-TEFb recruiters are also affected by PlaB treatment, we first performed ChIP-qPCR of RNAPII, BRD4, AFF4, MYC, and HTATSF1 in HeLa cells treated for 30 min with DMSO or PlaB on the promoters of *KPNB1, CCND2, LDLR*, and *SLCO4A1* (Figure 6B). While BRD4 and AFF4 recruitment is not affected on the four promoters after SF3B1 inhibition, we found that RNAPII, BRD4, and AFF4 are reduced from *KPNB1* region TSS +3.2, which represents productive transcription elongation, after PlaB treatment. For MYC and HTATSF1, we found a partial but not significant reduction in their recruitment to the TSS of these protein-coding genes. The results of western blots of BRD4, AFF4, and MYC on chromatin and nucleoplasm fractions of HeLa cells treated for 30 min with DMSO or PlaB (Figure 6C) indicate that only the association of AFF4 with chromatin is affected by PlaB treatment, with a reduction on chromatin coupled to an increase in the nucleoplasm. While we failed to ChIP another known P-TEFb recruiter, SRSF2, we found by whole cell extract western blot in HeLa cells that a 30 min treatment with PlaB reduces its total level (Figure 6D). To determine if SF3B1 inhibition results in an active degradation of SRSF2, we performed immunofluorescence experiments in HeLa cells treated with DMSO, the proteasome inhibitor Bortezomib, PlaB, and Bortezomib & PlaB (Figure 6E and F). In agreement with the western blot, PlaB treatment reduces SRSF2 fluorescent intensity but an 1 h pre-treatment of PlaB with Bortezomib blocks SRSF2 degradation. Interestingly, a short Bortezomib treatment alone also increases SRSF2 fluorescent intensity indicating that SRSF2 is actively turned over in the cell. These results indicate that in addition to SNW1, the P-TEFb recruiters AFF4 and SRSF2 are also affected by SF3B1 inhibition.

### Loss of P-TEFb from promoters after SF3B1 inhibition does not correlate with changes in RNAPII pausing

As multiple P-TEFb recruitment pathways are affected by SF3B1 inhibition, we investigated whether some genes were more affected by PlaB treatment. We first confirmed the previously observed loss of Cyclin T1 detected by spiked ChIP-seq on protein-coding genes after SF3B1 inhibition (Figure 7A and B and Supplementary Figure 7A and B) (30) with RNAPII and CDK9 ChIP-qPCR following 30 min treatment with 1 µM PlaB (Supplementary Figure 7C). SF3B1 inhibition promotes an increase in RNAPII pausing and a decrease in CDK9 recruitment on *KPNB1, SLCO4A1*, and *LDLR* promoters. As the degree of loss of CDK9 recruitment does not match the increase in RNAPII pausing on the three different promoters, we decided to investigate the global correlation between P-TEFb recruitment and RNAPII pausing using the previously-published spiked RNAPII and Cyclin T1 ChIP-seq (30) (Figure 7C and Supplementary Figure 7D-G). We found a positive correlation between P-TEFb level, RNAPII pausing index, and gene expression, with the genes possessing the highest Cyclin T1 level being on average associated with a higher RNAPII pausing index and higher gene expression. To understand better the effect of PlaB treatment on Cyclin T1 recruitment and RNAPII pausing, we separated promoters into four groups based on their level of Cyclin T1 ChIP-seq in the DMSO control condition and found that PlaB predominantly affects the Cyclin T1 level on genes associated with the highest amount of P-TEFb on the promoters (Figure 7D). Regarding the RNAPII pausing index, we observed that the increase in RNAPII pausing after PlaB treatment is strongest for genes that were moderately paused rather than genes with the lowest or highest pausing index (Figure 7E). We finally investigated whether the degree of loss of Cyclin T1 on promoters after PlaB treatment correlates with the extent of increase in RNAPII pausing. In agreement with CDK9 and RNAPII ChIP-qPCR results, the magnitude of increase of RNAPII pausing does not correlate with the extent of loss of Cyclin T1 (Figure 7F and G), with an increase in RNAPII pausing observed for genes associated with a clear loss of Cyclin T1 (Figure 7H) but also with a limited loss of Cyclin T1 (Supplementary Figure 7H). These results indicate that while SF3B1 inhibition affects P-TEFb recruitment and RNAPII pausing, there is no strict correlation between the two events and additional mechanisms are likely at play.

**Figure 7.**
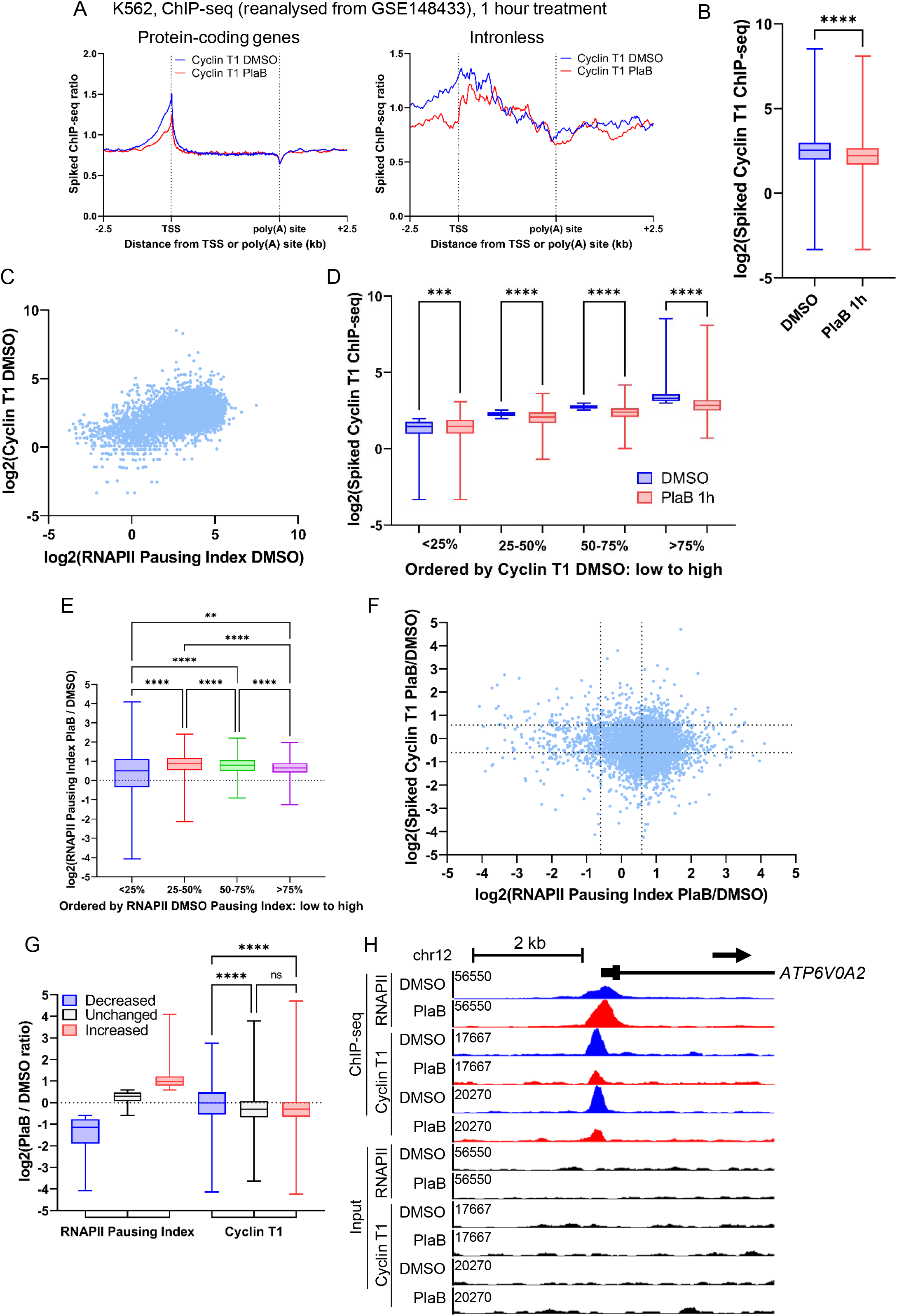
Loss of P-TEFb on promoters after SF3B1 inhibition does not correlate with changes in RNAPII pausing. (**A**) Metagene profile of spiked-in Cyclin T1 ChIP-seq in K562 cells treated with DMSO (blue) or 1 µM PlaB (red) for 60 min on scaled expressed protein-coding genes (left) and on intronless genes (right). (**B**) Boxplots (min to max, box showing 25%, median, and 75%) of spiked-in Cyclin T1 ChIP-seq on promoter regions of protein-coding genes, n = 6,960. Statistical test: Wilcoxon matched-pairs signed rank test. P-value: ****: < 0.0001. (**C**) XY plot showing in K562 cells the log2 of the RNAPII DMSO pausing index versus the log2 of the spiked-in Cyclin T1 DMSO ChIP-seq signal across promoters, n = 6,960. (**D**) Boxplots (min to max, box showing 25%, median, and 75%) of spiked-in Cyclin T1 ChIP-seq after DMSO and PlaB treatments on promoter regions of protein-coding genes that have been categorised based on their Cyclin T1 DMSO signal from low (<25%) to high (>75%). Statistical test: Kruskal-Wallis test, n = 1,740 promoter regions per category. P-value: *** < 0.001, **** < 0.0001. (**E**) Boxplots (min to max, box showing 25%, median, and 75%) of the log2 of the RNAPII pausing index PlaB / DMSO ratio of protein-coding genes that have been categorised based on their DMSO pausing index from low pausing (<25%) to high pausing (>75%). Statistical test: Kruskal-Wallis test, n = 1,740 promoter regions per category. P-value: ** < 0.01, **** < 0.0001. (**F**) XY plot showing in K562 cells the log2 of the RNAPII pausing index PlaB / DMSO ratio versus the log2 of the spiked-in Cyclin T1 PlaB / DMSO ratio on the promoter regions of these protein-coding genes, n = 6,960. (**G**) Boxplots (min to max, box showing 25%, median, and 75%) of the log2 of the PlaB / DMSO ratio for RNAPII pausing index and Cyclin T1 signal on promoter regions for protein-coding genes that have been categorised based on their changes in RNAPII pausing index. Decreased: PlaB / DMSO ratio ≤ 0.667 (n = 387), Unchanged: PlaB / DMSO ratio between 0.667 and 1.5 (n = 2,464), Increased: PlaB / DMSO ratio ≥ 1.5 (n = 4,505). Statistical test: Kruskal-Wallis test. P-value: ns: not significant, **** < 0.0001. (**H**) Screenshot of the genome browser for K562 spiked-in total RNAPII and Cyclin T1 ChIP-seq DMSO (blue) and PlaB (red) and their respective Input (black) tracks around the promoter region of the protein-coding gene *ATP6V0A2*. The arrow indicates the sense of transcription.

## DISCUSSION

While it was previously shown that inhibition of SF3B1 results in loss of Cyclin T1 from chromatin, the mechanism was unknown (30). HTATSF1, which plays roles in transcription elongation and splicing, was proposed as a key mediator between SF3B1 and P-TEFb (30). Here we show that SF3B1 is in complex with P-TEFb, HTATSF1, and SNW1 on chromatin. In addition to limited export of P-TEFb from the nucleus, PlaB treatment also promotes the export of chromatin-associated SNW1 to the cytoplasm. We show that P-TEFb phosphorylates several residues of SF3B1 and that an additional transcriptional kinase, CDK12, regulates SF3B1 Thr313 phosphorylation. Furthermore, we demonstrate that PlaB affects the chromatin association of two other known P-TEFb recruiters, SRSF2 and AFF4. Altogether, these results indicate that SF3B1 inhibition affects P-TEFb recruitment to chromatin via multiple pathways (Figure 8) and that surprisingly, SF3B1 activity is required to maintain P-TEFb levels in the nucleus.

**Figure 8.**
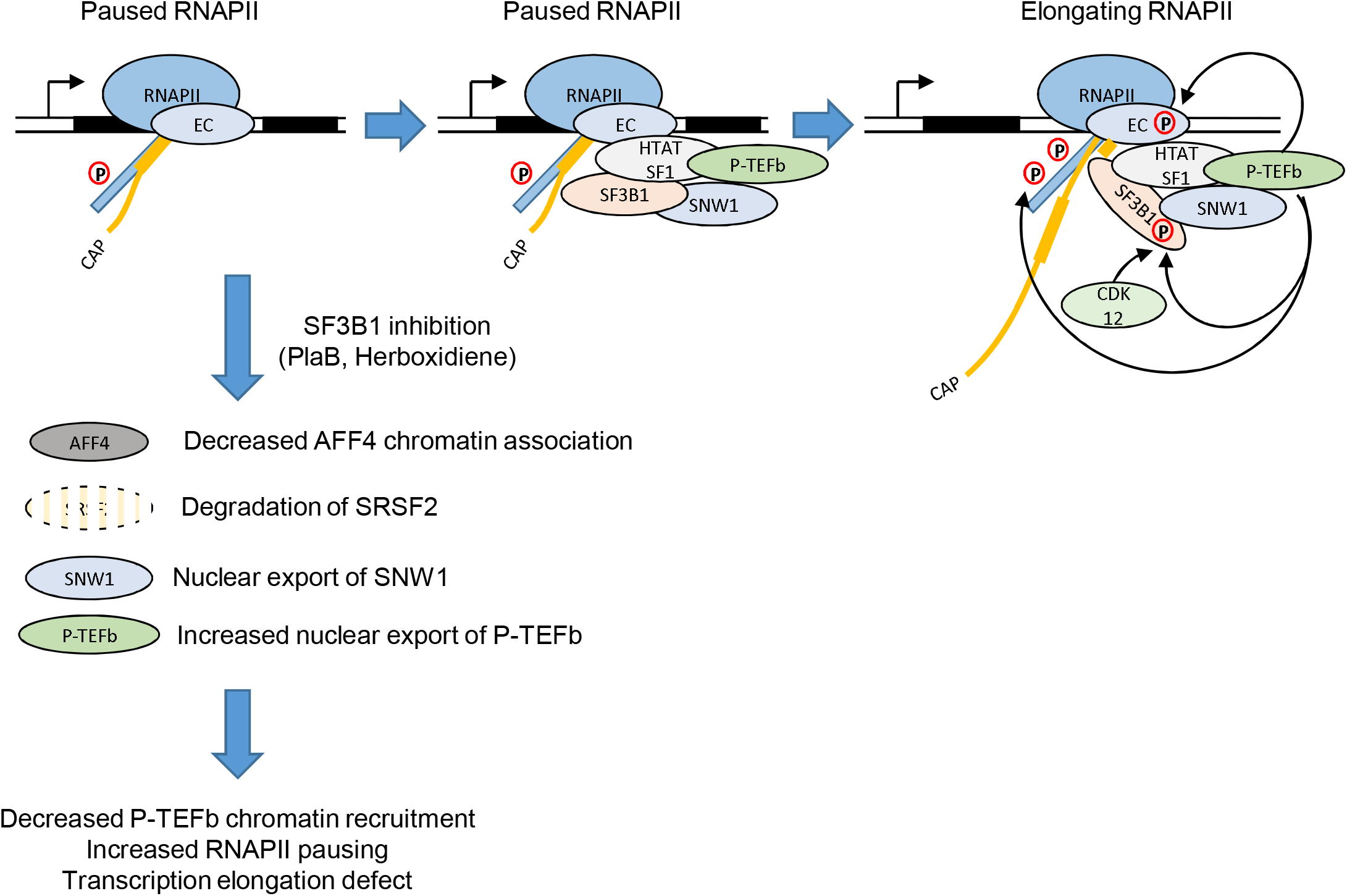
Multiple P-TEFb recruitment pathways to chromatin are affected by SF3B1 inhibition. Schematic showing how SF3B1/HTATSF1/SNW1 could recruit P-TEFb for RNAPII pause release and phosphorylation of SF3B1 during pre-mRNA splicing. The schematic also shows P-TEFb recruitment pathways to chromatin affected by SF3B1 inhibition. EC: elongation complex containing NELF (Paused RNAPII step) and DSIF. CDK12 recruitment/activity may also be affected by SF3B1 inhibition.

More specifically, we confirmed that P-TEFb interacts with HTATSF1 (17,18), SNW1 (21,22), and SF3B1 (75), and that SF3B1 interacts with HTATSF1 (78,79) and SNW1 (80), and found that most of these interactions occur on chromatin only. SF3B1 and HTATSF1 instead also interact in the nucleoplasm and cytoplasm. However, a major question remains-do the interactions between P-TEFb and SF3B1/HTATSF1/SNW1 bring P-TEFb to the paused RNAPII to promote transcription elongation and/or do they bring P-TEFb to phosphorylate SF3B1 to regulate pre-mRNA splicing?

In support of its role in recruitment of P-TEFb to paused RNAPII, we found by ChIP-qPCR that HTATSF1 is present at TSSs, in agreement with previous results (81). Additionally, HTATSF1 is known to interact with SPT5 (82), a known target of P-TEFb kinase activity (83).

Unfortunately, we failed to ChIP SF3B1 or SNW1 using commercial antibodies or a FLAG-tagged SF3B1 and re-analysis of previously published SF3B1 ChIP-seq datasets (39,54) indicates limited enrichment of SF3B1 over background across protein-coding genes (Supplementary Figure 7I). SNW1 was previously shown to interact with P-TEFb to promote expression of NF-κB target genes following TNFα treatment (22). However, SNW1 knockdown does not reduce CDK9 and RNAPII recruitment to the promoters of NF-κB target genes after TNFα treatment (22), questioning the role of SNW1 in directly recruiting P-TEFb to paused RNAPII. Based on these results, HTATSF1 rather than SNW1 could therefore be directly involved in the recruitment of P-TEFb for RNAPII pause release. However, HTATSF1 alone is clearly unable to compensate for the effect of SF3B1 inhibition, indicating that P-TEFb recruitment via HTATSF1 alone is insufficient.

Expanding on our previous observation that CDK9 phosphorylates SF3B1 on Thr142 (38), we show here that CDK9 also phosphorylates Ser129 and to a lesser extent Thr211 residues whereas CDK12 phosphorylates Thr313. However, while a role of CDK9 in splicing has been proposed (84) and P-TEFb can bind the 5’ splice sites of mRNAs (72), P-TEFb’s role in splicing remains poorly understood as the rapid loss of transcription following CDK9 inhibition hinders analysis of co-transcriptional splicing (38). In the case of CDK12, its inhibition with THZ531 (85), a CDK12/CDK13 inhibitor (86), or in an CDK12 analogue-sensitive kinase cell line (87) was found to affect co-transcriptional splicing by controlling the association of SF3B1 with RNAPII (85). As CDK12 inhibition promotes a transcription elongation defect (43,56,88,89), it cannot be ruled out that this could also influence co-transcriptional splicing as an optimal RNAPII elongation rate facilitates splicing (90).

In addition to the nuclear export of SNW1 following PlaB treatment, SF3B1 inhibition also affects AFF4 and SRSF2, two other known recruiters of P-TEFb. AFF4 is part of the SEC that promotes transcription elongation (14). Loss of AFF4, and more generally of the SEC, from chromatin would therefore reduce the amount of elongating RNAPII. In addition, loss of AFF4 likely exacerbates the loss of P-TEFb from chromatin. Surprisingly, we found that PlaB treatment results in the active degradation of SRSF2 by the nuclear proteasome. It has been previously shown that SRSF2 acetylation by KAT5 (TIP60) promotes its degradation (91), which could also be the case following PlaB treatment.

Our experiments trying to block the nuclear export of P-TEFb with LMB did not work, likely due to a too short pre-treatment time of 1 h before PlaB treatment. As LMB affects the nuclear import of CRM1, CRM1 can continue to shuttle protein out of the nucleus until there is no CRM1 available in the nucleus (92). Therefore, a longer pre-treatment time with LMB would be required before treating with PlaB. However, we demonstrate that LMB itself affects transcription after 1 h, which to our knowledge has not been previously shown and should therefore be kept in mind when interpreting results obtained with LMB.

SF3B1 inhibition was already known to decrease transcription elongation of intron-containing and intronless genes (30,34) but we also found that transcription termination, but not transcription elongation, of histone genes is affected by PlaB treatment. Interestingly, the effect of SF3B1 inhibition on termination of transcription of histone genes is similar to CDK9 inhibition, which causes premature termination of transcription of protein-coding genes (38,70,93). As SF3B1 is found on histone genes (Supplementary Figure 7I) and PlaB treatment reduces Cyclin T1 recruitment to histone genes (Supplementary Figure 3A), the effect of SF3B1 inhibition on transcription termination of histone genes is likely mediated by the loss of P-TEFb. It remains unclear whether this premature termination of transcription of histone genes affects 3’end processing of histone transcripts as CDK9 inhibition was previously found not to decrease 3’end processing of histone H2B gene transcripts (94).

A surprising observation is that while Cyclin T1 recruitment to the TSS region of protein-coding genes is positively correlated to RNAPII pausing and RNA expression level, there is no correlation between the extent of Cyclin T1 loss and the magnitude of increase of the RNAPII pausing index following PlaB treatment. These results indicate that changes in P-TEFb recruitment are not sufficient by themselves to explain increased RNAPII pausing and that therefore additional mechanisms regulating transcription initiation, RNAPII pausing, and/or RNAPII premature termination might also be affected by SF3B1 inhibition.

In conclusion, we have shown that SF3B1 inhibition causes a loss of recruitment of P-TEFb to chromatin via multiple pathways. However, P-TEFb loss does not entirely explain the increase in RNAPII pausing and decrease in RNAPII elongation following PlaB treatment. It remains to be determined what additional mechanisms are affected by SF3B1 inhibition to understand the role of SF3B1, and more generally of splicing factors, in the regulation of transcription.

## DATA AVAILABILITY

mNET-seq data have been deposited in GEO under accession number GSE221279.

## SUPPLEMENTARY DATA

Supplementary Data are available at NAR online.

## AUTHOR CONTRIBUTIONS

Gilbert Ansa: Formal analysis, Investigation, Methodology, Validation, Writing—review & editing. Shona Murphy: Funding acquisition, Conceptualization, Supervision, Writing—review & editing. Michael Tellier: Conceptualization, Formal analysis, Investigation, Methodology, Supervision, Validation, Visualization, Writing—original draft, Writing—review & editing.

## ACKNOWLEDGEMENTS

We thank the High-Throughput Genomics Group at the Wellcome Trust Centre for Human Genetics for sequencing.

## FUNDING

This work was supported by the Wellcome Trust [WT106134AIA and WT210641/Z/18/Z to S.M.]. Funding for open access charge: University of Leicester.

## CONFLICT OF INTEREST

Conflict of interest statement. None declared.

## Notes

### Competing Interest Statement

The authors have declared no competing interest.

