## Supplementary material for "Inhibition of SF3B1 affects recruitment of P-TEFb to chromatin through multiple mechanisms": Supp FI

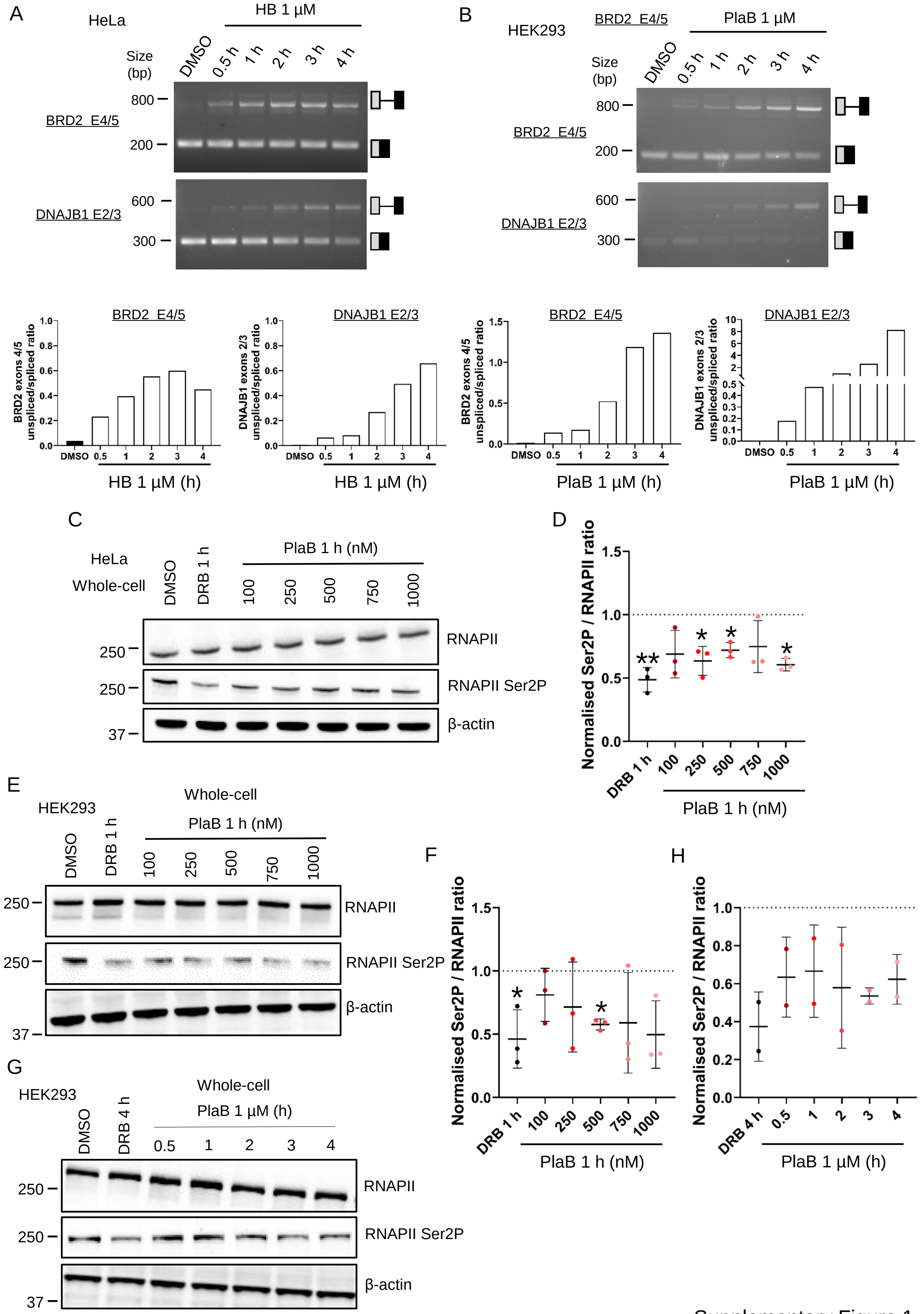


Supplementary Figure 1

**Supplementary Figure 1. Rapid inhibition of SF3B1 decreases pre-mRNA splicing and RNAPII transcriptional activity.**

(**A**) RT-PCR with primers amplifying the intron located between exons 4 and 5 of BRD2 or exons 2 and 3 of DNAJB1. HeLa cells were treated with DMSO or 1 µM HB for 30 min to 4 h. The location of the spliced and unspliced RNA is shown on the left of the panel. The percentages of unspliced RNA compared to total (spliced + unspliced) are shown below. (**B**) RT-PCR with primers amplifying the intron located between exons 4 and 5 of BRD2 or exons 2 and 3 of DNAJB1. HEK293 cells were treated with DMSO or 1 µM HB for 30 min to 4 h. The location of the spliced and unspliced RNA is shown on the left of the panel. The percentages of unspliced RNA compared to total (spliced + unspliced) are shown below. (**C**) Representative whole cell western blots of total RNAPII, RNAPII Ser2 phosphorylation, and β-actin (loading control) from HeLa cells treated with DMSO, 100 µM DRB for 1 h, or 100 - 1000 nM PlaB for 1 h. (**D**) Quantification of the western blots shown in (C) presented as RNAPII Ser2P / total RNAPII following normalisation to the loading control and to the DMSO condition. Statistical test: Kruskal-Wallis test, n = 3 biological replicates. P-value: * < 0.05, ** < 0.01. (**E**) Representative whole cell western blots of total RNAPII, RNAPII Ser2 phosphorylation, and β-actin (loading control) from HEK293 cells treated with DMSO, 100 µM DRB for 1 h, or 100 - 1000 nM PlaB for 1 h. (**F**) Quantification of the western blots shown in (E) presented as RNAPII Ser2P/total RNAPII following normalisation to the loading control and to the DMSO condition. Statistical test: Kruskal-Wallis test, n = 3 biological replicates. P-value: * < 0.05. (**G**) Representative whole cell western blots of total RNAPII, RNAPII Ser2 phosphorylation, and β-actin (loading control) from HEK293 cells treated with DMSO, 100 µM DRB for 4 h, or 1 µM PlaB for 30 min to 4 h. (**H**) Quantification of the western blots shown in (G) presented as RNAPII Ser2P / total RNAPII following normalisation to the loading control and to the DMSO condition, n= 2 biological replicates.

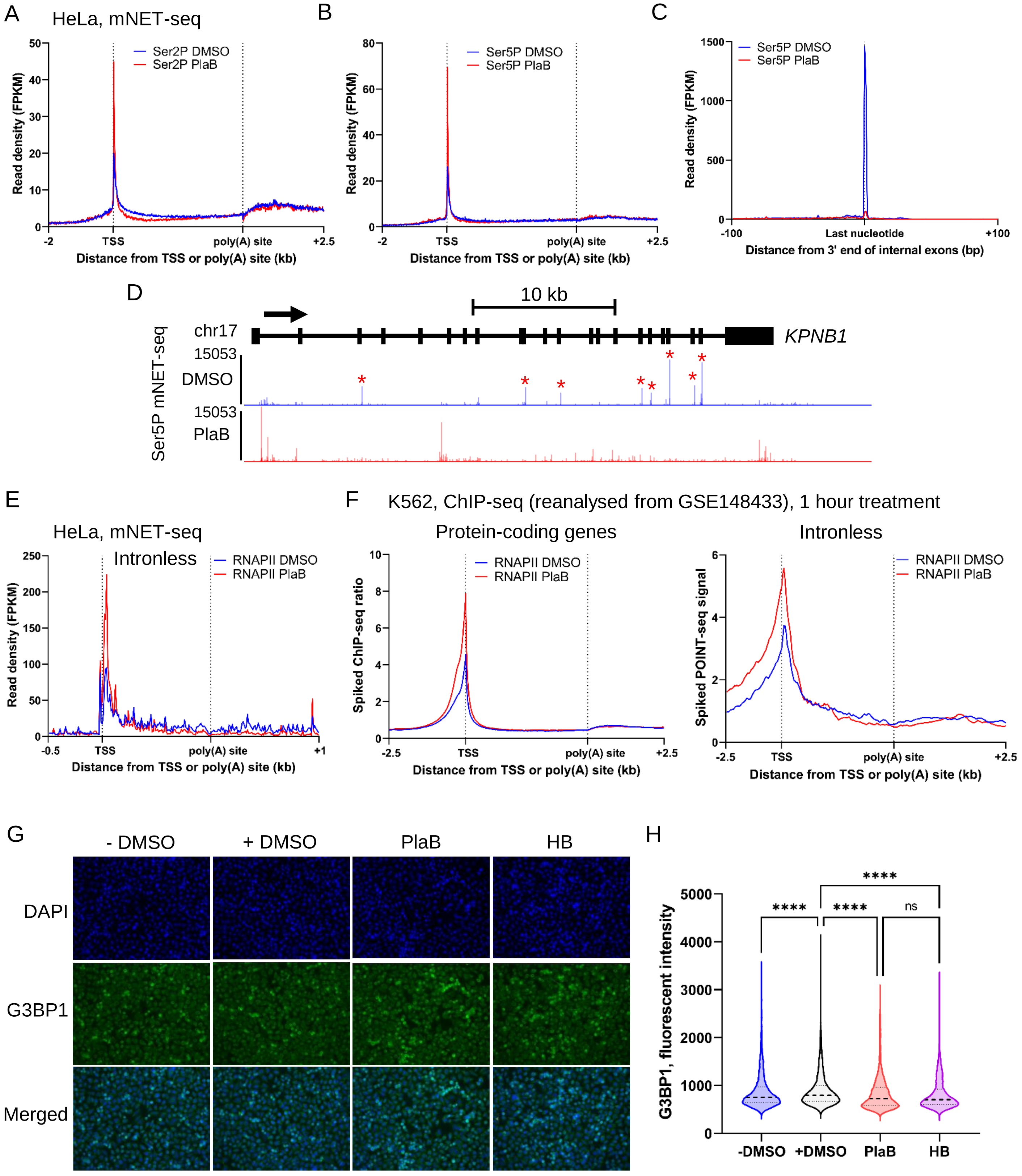


Supplementary Figure 2

**Supplementary Figure 2. SF3B1 inhibition increases RNAPII pausing.**

(**A**) Metagene profile of RNAPII Ser2P mNET-seq in HeLa cells treated with DMSO (blue) or 1 µM PlaB (red) for 30 min on scaled expressed protein-coding genes. (**B**) Metagene profile of RNAPII Ser5P mNET-seq in HeLa cells treated with DMSO (blue) or 1 µM PlaB (red) for 30 min on scaled expressed protein-coding genes. (**C**) Metagene profile of RNAPII Ser5P mNET-seq in HeLa cells treated with DMSO (blue) or 1 µM PlaB (red) for 30 min around the 5' splice site (5'SS) of internal exons, i.e., neither first or last exons. (**D**) Screenshot of the genome browser of RNAPII Ser5P mNET-seq DMSO (blue) and PlaB (red) tracks of the protein-coding gene *KPNB1*. Red stars indicate the location of co-transcriptional splicing events. The arrow indicates the sense of transcription. (**E**) Metagene profile of total RNAPII mNET-seq in HeLa cells treated with DMSO (blue) or 1 µM PlaB (red) for 30 min on scaled expressed intronless genes. (**F**) Metagene profile of spiked-in total RNAPII ChIP-seq in K562 cells treated with DMSO (blue) or 1 µM PlaB (red) for 60 min on scaled expressed protein-coding genes (left) or intronless genes (right). (**G**) Representative images of immunofluorescence analysis of G3BP1 in HeLa cells treated with nothing, DMSO, 1 µM PlaB, or 1 µM HB for 30 min. G3BP1 (green), DAPI (blue), scale bars: 50 µm. (**H**) Quantification of G3BP1 fluorescent intensity per nucleus for nothing (blue), DMSO (black), PlaB (red), and HB (purple). Boxplot settings are: min to max values with the box showing 25-75 percentile range. 10,522 nuclei were quantified per condition. Statistical test: Kruskal-Wallis test. P-value: ns: not significant, **** < 0.0001.

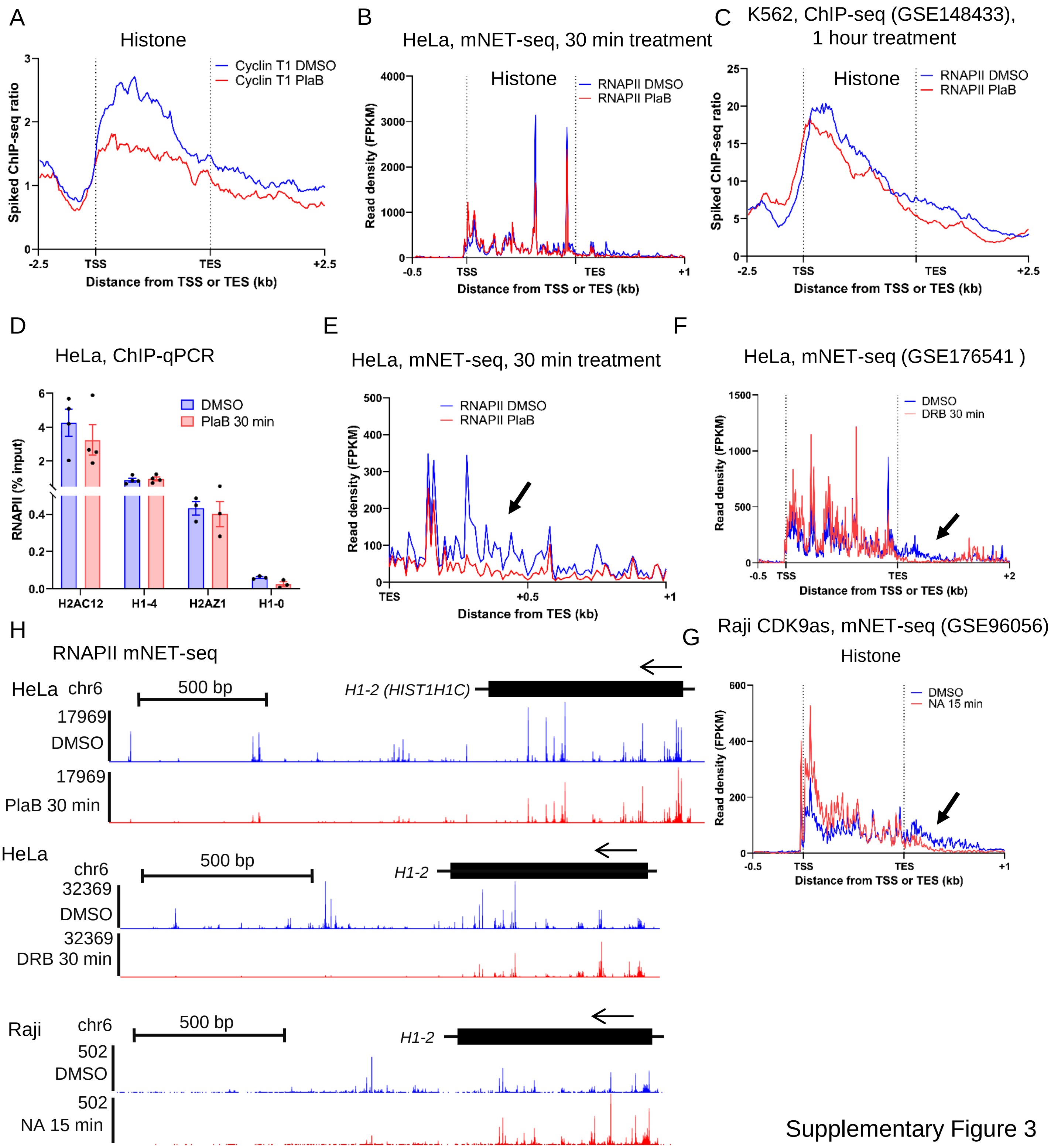


Supplementary Figure 3

**Supplementary Figure 3. SF3B1 or CDK9 inhibition decreases transcription downstream of histone gene bodies.**

(**A**) Metagene profile of spiked-in Cyclin T1 ChIP-seq in K562 cells treated with DMSO (blue) or 1 µM PlaB (red) for 60 min on scaled expressed histone genes. (**B**) Metagene profile of total RNAPII mNET-seq in HeLa cells treated with DMSO (blue) or 1 µM PlaB (red) for 30 min on scaled expressed histone genes. (**C**) Metagene profile of spiked-in total RNAPII ChIP-seq in K562 cells treated with DMSO (blue) or 1 µM PlaB (red) for 60 min on scaled expressed histone genes. (**D**) Total RNAPII ChIP-qPCR on the gene body of *H2AC12*, *H1-4*, *H2AZ1*, and *H1-0* in HeLa cells treated with DMSO or 1 µM PlaB for 30 min, n = 3-4 biological replicates. (**E**) Zoom on the metagene profile of total RNAPII mNET-seq in HeLa cells treated with DMSO (blue) or 1 µM PlaB (red) for 30 min on the transcription termination region of histone genes. (**F**) Metagene profile of total RNAPII mNET-seq in HeLa cells treated with DMSO (blue) or 100 µM DRB (red) for 30 min on scaled expressed histone genes. (**G**) Metagene profile of total RNAPII mNET-seq in Raji CDK9 analogue-sensitive cells treated with DMSO (blue) or 5 µM 1-NA-PP1 (red) for 15 min on scaled expressed histone genes. (**H**) Screenshot of the genome browser of total RNAPII mNET-seq in HeLa cells treated with DMSO (blue), 1 µM PlaB (red), or 100 µM DRB (red) for 30 min or of total RNAPII mNET-seq in Raji CDK9 analogue-sensitive cells treated with DMSO (blue) or 5 µM 1-NA-PP1 (red) for 15 min. Tracks are for the histone gene *H1-2*. The arrow indicates the sense of transcription.

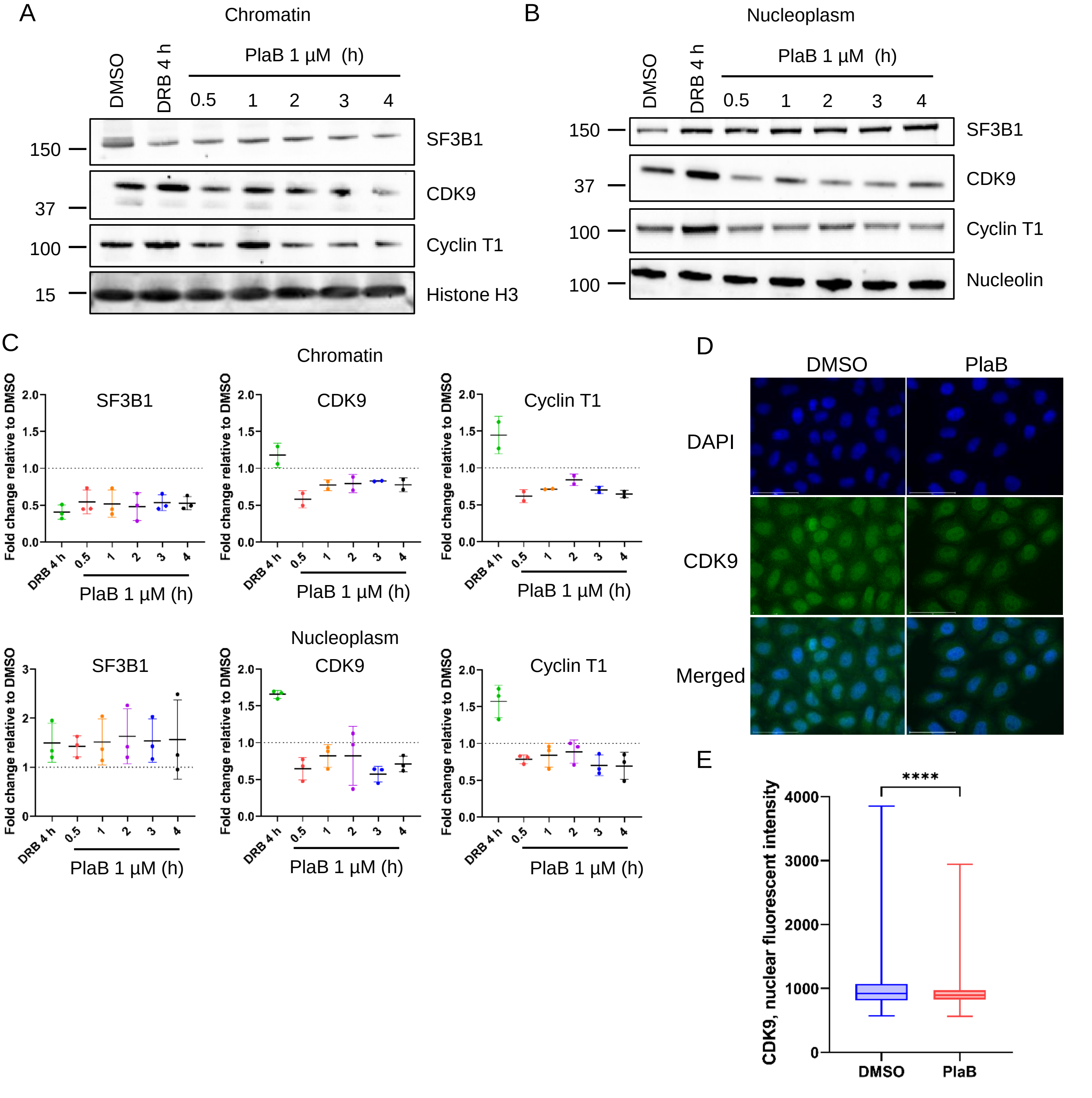


Supplementary Figure 4

**Supplementary Figure 4. SF3B1 inhibition increases P-TEFb nuclear export.**

(**A**) Representative western blots of SF3B1, CDK9, Cyclin T1, and histone H3 (loading control) from chromatin fraction of HeLa cells treated with DMSO, 100 µM DRB for 4 h, and 1 µM PlaB for 30 min to 4 h. (**B**) Representative western blots of SF3B1, CDK9, Cyclin T1, and nucleolin (loading control) from nucleoplasm fraction of HeLa cells treated with DMSO, 100 µM DRB for 4 h, and 1 µM PlaB for 30 min to 4 h. (**C**) Quantification of western blots of SF3B1, CDK9, and Cyclin T1 from HeLa chromatin and nucleoplasm fractions following treatment with 100 µM DRB for 4 h, and 1 µM PlaB for 30 min to 4 h. The quantifications have been normalised to loading control and to DMSO, n = 2-3 biological replicates. (**D**) Representative images of immunofluorescence analysis of CDK9 in HeLa cells treated with DMSO or 1 µM PlaB for 30 min. CDK9 (green), DAPI (blue), scale bars: 50 µm. (**E**) Quantification of CDK9 fluorescent intensity per nucleus for DMSO (blue) and PlaB (red). Boxplot settings are: min to max values with the box showing 25-75 percentile range. 3,478 nuclei were quantified per condition. Statistical test: Kruskal-Wallis test. P-value: **** < 0.0001.

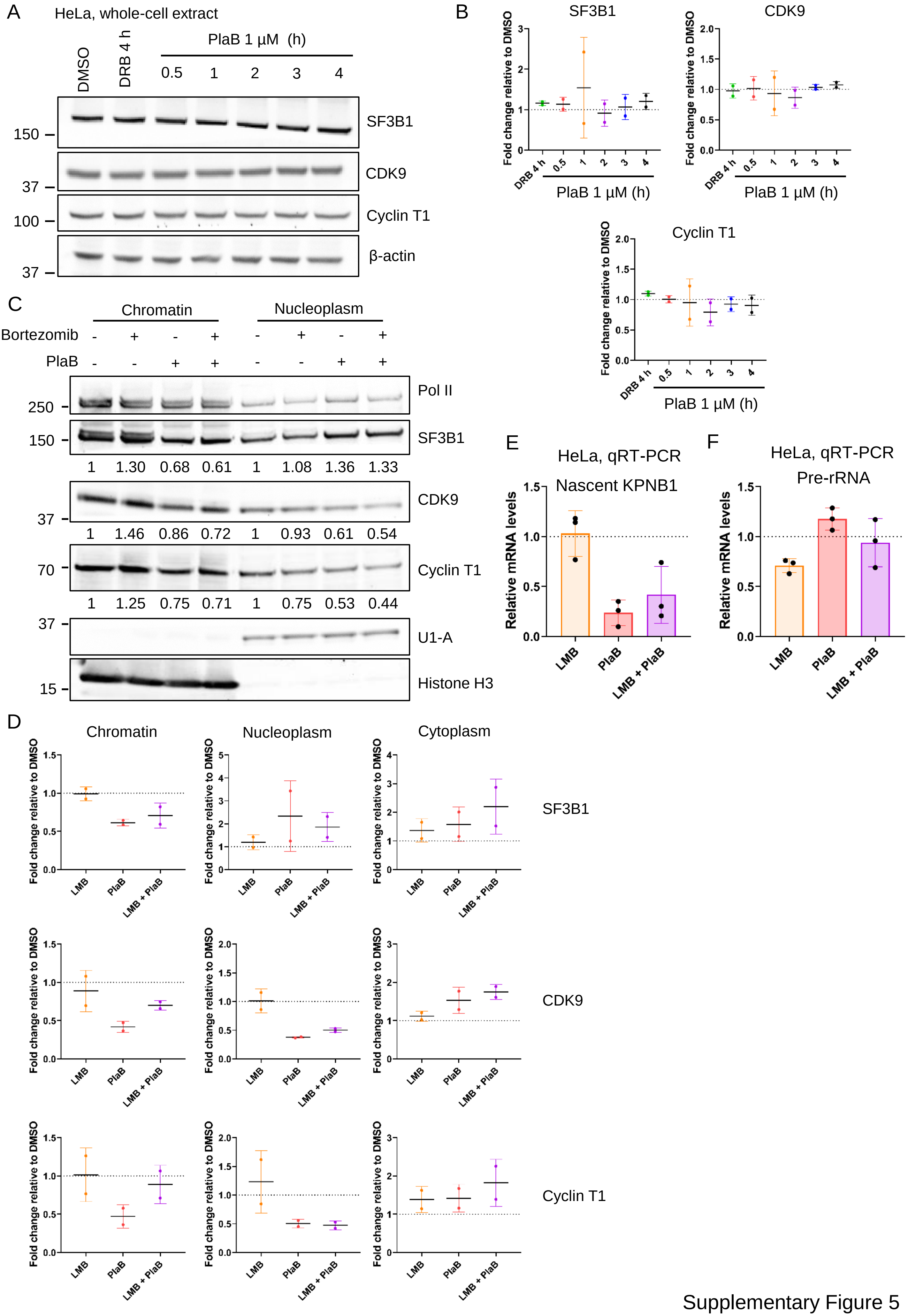


Supplementary Figure 5

**Supplementary Figure 5. SF3B1 inhibition does not promote degradation of SF3B1 and P-TEFb.**

(**A**) Representative western blots of SF3B1, CDK9, Cyclin T1, and β-actin (loading control) from whole cell extract of HeLa cells treated with DMSO, 100 µM DRB for 4 h, and 1 µM PlaB for 30 min to 4 h. (**B**) Quantification of western blots of SF3B1, CDK9, and Cyclin T1 from HeLa whole cell extract following treatment with 100 µM DRB for 4 h, and 1 µM PlaB for 30 min to 4 h. The quantifications have been normalised to loading control and to DMSO, n = 2 biological replicates. (**C**) Western blots of total RNAPII, SF3B1, CDK9, Cyclin T1, U1-A (loading control), and histone H3 (loading control) from chromatin and nucleoplasm fractions of HeLa cells treated with DMSO, 1 µM Bortezomib, or 1 µM PlaB. Treatments were performed as follow: DMSO 1h/DMSO 30 min (-/-), Bortezomib 1h/DMSO 30 min (+/-), DMSO 1h/PlaB 30 min (-/+), Bortezomib 1h/PlaB 30 min (+/+). (**D**) Quantification of western blots of SF3B1, CDK9, and Cyclin T1 from chromatin, nucleoplasm, and cytoplasm fractions of HeLa cells treated with DMSO, ethanol, 0.02 µM Leptomycin B, or 1 µM PlaB. Treatments were performed as follow: Ethanol 1h/DMSO 30 min (DMSO), LMB 1h/DMSO 30 min (LMB), Ethanol 1h/PlaB 30 min (PlaB), LMB 1h/PlaB 30 min (LMB + PlaB). (**E**) qRT-PCR of nascent KPNB1 transcript in HeLa cells treated with DMSO, LMB, PlaB, or LMB & PlaB, n = 3 biological replicates. (**F**) qRT-PCR of pre-rRNA transcript in HeLa cells treated with DMSO, LMB, PlaB, or LMB & PlaB, n = 3 biological replicates.

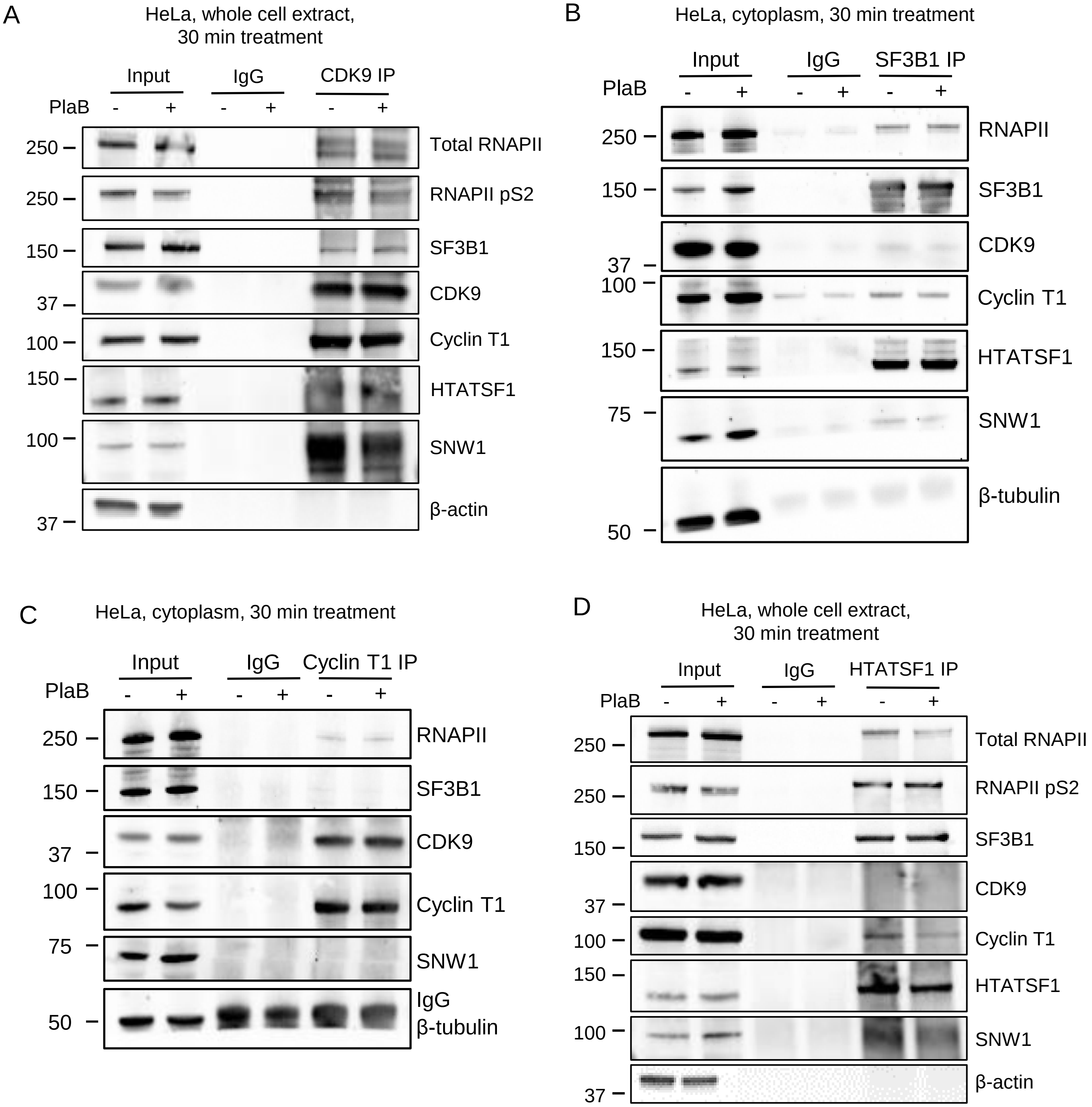


Supplementary Figure 6

**Supplementary Figure 6. SF3B1 inhibition affects predominantly SNW1 in the chromatin-associated SF3B1/HTATSF1/SNW1/P-TEFb complex.**

(**A**) Co-immunoprecipitation of CDK9 from whole cell extract of HeLa cells treated with DMSO or 1 µM PlaB for 30 min followed by western blots of total RNAPII, RNAPII Ser2 phosphorylation, SF3B1, CDK9, Cyclin T1, HTATSF1, SNW1, and β-actin (negative control). (**B**) Co-immunoprecipitation of SF3B1 from cytoplasm of HeLa cells treated with DMSO or 1 µM PlaB for 30 min followed by western blots of RNAPII, SF3B1, CDK9, Cyclin T1, HTATSF1, SNW1, and β-tubulin (negative control). (**C**) Co-immunoprecipitation of Cyclin T1 from cytoplasm of HeLa cells treated with DMSO or 1 µM PlaB for 30 min followed by western blots of RNAPII, SF3B1, CDK9, Cyclin T1, SNW1, and β-tubulin (negative control). (**D**) Co-immunoprecipitation of HTATSF1 from whole cell extract of HeLa cells treated with DMSO or 1 µM PlaB for 30 min followed by western blots of total RNAPII, RNAPII Ser2 phosphorylation, SF3B1, CDK9, Cyclin T1, HTATSF1, SNW1, and β-actin (negative control).

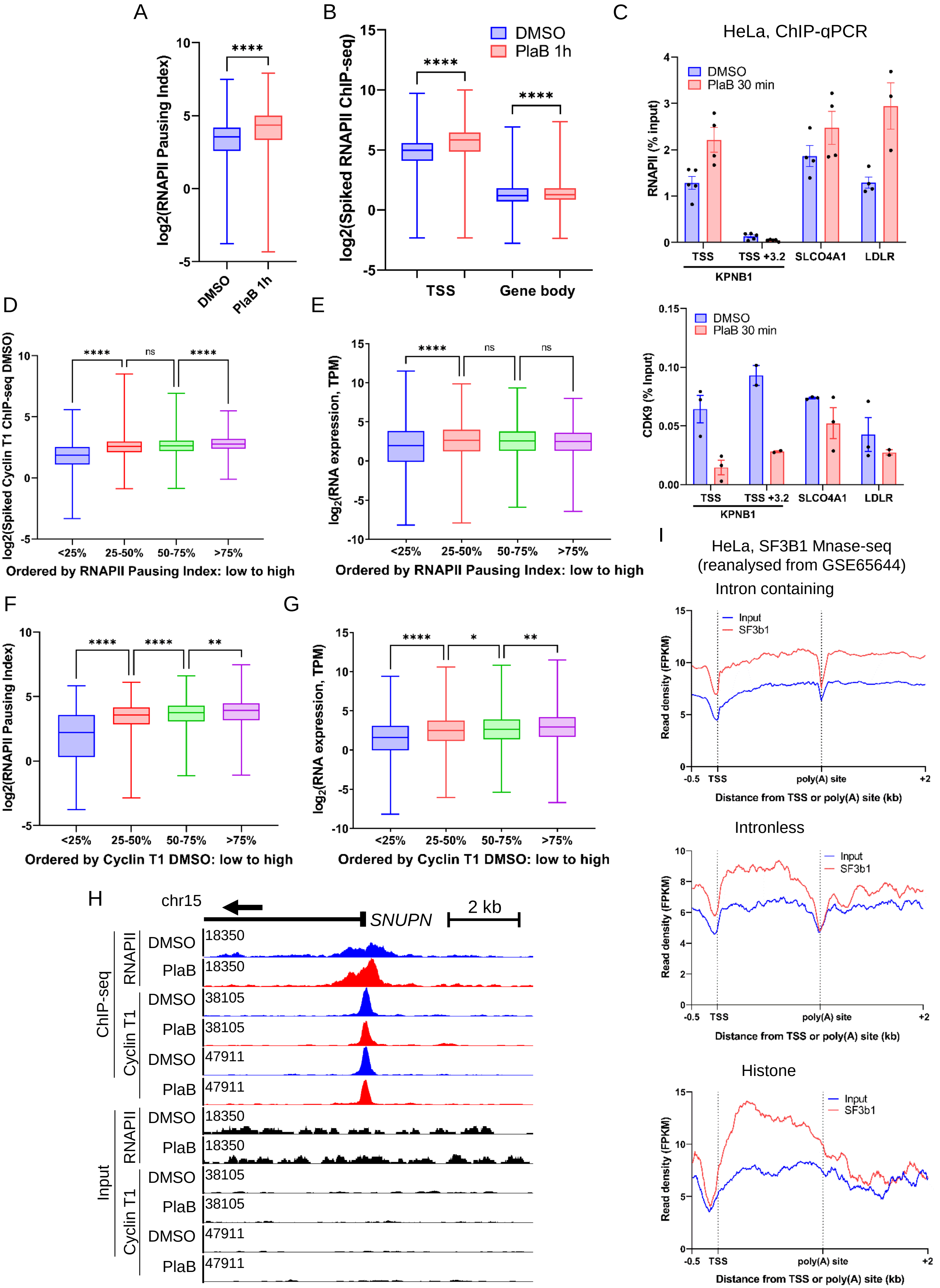


Supplementary Figure 7

**Supplementary Figure 7. Loss of P-TEFb on promoters after SF3B1 inhibition does not correlate with changes in RNAPII pausing.**

(**A**) Boxplots (min to max, box showing 25%, median, and 75%) of RNAPII pausing index in K562 cells treated with DMSO or 1 µM PlaB for 1 h, n = 6,960. Statistical test: Wilcoxon matched-pairs signed rank test. P-value: ****: < 0.0001. (**B**) Boxplots (min to max, box showing 25%, median, and 75%) of spiked-in total RNAPII ChIP-seq signal in K562 cells treated with DMSO or 1 µM PlaB for 1 h on promoter regions and across the gene body, n = 6,960. Statistical test: Wilcoxon matched-pairs signed rank test. P-value: ****: < 0.0001. (**C**) Total RNAPII and CDK9 ChIP-qPCR on the promoter regions of *KPNB1*, *SLCO4A1*, and *LDLR* in HeLa cells treated with DMSO or 1 µM PlaB for 30 min, n = 2-4 biological replicates. The RNAPII ChIP-qPCR data are the same as the ones shown in Figure 6B (same ChIP-qPCR experiments). (**D**) Boxplots (min to max, box showing 25%, median, and 75%) of the log2 of spiked-in Cyclin T1 DMSO ChIP-seq on promoter regions of protein-coding genes that have been categorised based on their DMSO RNAPII pausing index from low pausing (<25%) to high pausing (>75%). Statistical test: Kruskal-Wallis test, n = 1,740 per category. P-value: ns: not significant, **** < 0.0001. (**E**) Boxplots (min to max, box showing 25%, median, and 75%) of the log2 of RNA-seq expression level (TPM) of protein-coding genes that have been categorised based on their DMSO pausing index from low pausing (<25%) to high pausing (>75%). Statistical test: Kruskal-Wallis test, n = 1,740 per category. P-value: ns: not significant, **** < 0.0001. (**F**) Boxplots (min to max, box showing 25%, median, and 75%) of the log2 of DMSO RNAPII pausing index of protein-coding genes that have been categorised based on their Cyclin T1 DMSO signal from low (<25%) to high (>75%) on promoter regions. Statistical test: Kruskal-Wallis test, n = 1,740 per category. P-value: ** < 0.01, **** < 0.0001. (**G**) Boxplots (min to max, box showing 25%, median, and 75%) of RNA-seq expression level (TPM) of protein-coding genes that have been categorised based on their Cyclin T1 DMSO signal from low (<25%) to high (>75%) on promoter regions. Statistical test: Kruskal-Wallis test, n = 1,740 per category. P-value: * < 0.05, ** < 0.01, **** < 0.0001. (**H**) Screenshot of the genome browser for K562 spiked-in total RNAPII and Cyclin T1 ChIP-seq DMSO (blue) and PlaB (red) and their respective Input (black) tracks around the promoter region of the protein-coding gene *SNUPN*. The arrow indicates the sense of transcription. (**I**) Metagene profile of SF3B1 MNase-seq (red) and its Input (blue) in untreated HeLa cells on scaled expressed intron-containing, intronless, and histone genes.
